# Concordant developmental expression profiles of orthologues in highly divergent Bilateria

**DOI:** 10.1101/786723

**Authors:** Luca Ferretti, Andrea Krämer-Eis, Philipp H. Schiffer

## Abstract

Bilateria are the predominant clade of animals on earth. Despite having evolved a large variety of body-plans and developmental modes, they are characterized by common morphological traits. However, it is not clear if clade-specific genes can be linked to these traits, distinguishing bilaterians from non-bilaterians, with their less complex body morphology. Comparing proteomes of bilaterian and non-bilaterian species in an elaborate computational pipeline we aimed to find and define a set of of bilaterian-specific genes. Finding no high-confidence set of such genes, we nevertheless detected an evolutionary signal possibly uniting the highly diverse bilaterian taxa. Using a novel multi-species GO-enrichment method, we determined the functional repertoire of genes that are widely conserved among Bilateria. We found that these genes contribute to morphogenesis, neuronal-system and muscle development, processes that have been described as different between bilaterians and non-bilaterians. Analyzing gene expression profiles in three very distantly related bilaterina species, we find characteristic peaks at comparable stages of development and a delayed onset of expression in embryos. In particular, the expression of the conserved genes appears to peak at the phylotypic stage of different bilaterian phyla. In summary, our data underpin the orthologue conjecture and illustrate how development connects distantly related Bilateria after millions of years of divergence, pointing to processes potentially separating them from non-bilaterians.

## 1. Introduction

Bilateria represent by far the largest monophyletic group in the animal kingdom. They comprise about 99% of the extant eumetazoans [1] and are classified into 32 phyla [2]. The taxon ‘Bilateria’ has been defined on the basis of morphological key innovations, namely bilateral symmetry, triploblasty, an enhanced nervous system and a complex set of cell types [3]. Probably the most striking observation is the existence of a large variety of body plans, accompanied by high morphological diversity in many bilaterian phyla [4].

It is thought that the major bilaterian phyla emerged in a fast radiation during the early Cambrian, about 540 million years ago [1], but the appearance of the common bilaterian ancestor, and its exact timing, is controversial [5, 6, 7, 8]. It is also difficult to define unifying morphological properties for larval or adult Bilateria, as their descendants underwent extensive re-modellings (including secondary reductions and simplifications) of body forms in the lineages leading to extant crown clades. Nevertheless, the general process of development from single-cell to adult is conserved among all animals. Recently, evidence for the hypothesis that the developmental transcriptome might be a conserved trait across diverse groups of animals has been found [9, 10]. In particular, several studies described a conserved phylotypic period mid development based on transcriptomic analyses [11, 12, 13], akin to the morphological hourglass model of developmental progression [14, 15], itself an extension of von Baers reverse funnel model of development in animals [16].

Despite these similarities in the global transcriptome, comparisons at the molecular level reveal that many developmentally important genes are older than the ancestor of Bilateria. Among them are determinants of dorso-ventral and anterior-posterior patterning, eye formation, segmentation, and heart development [5, 17, 18, 19, 20]. In a recent study, it was estimated that up to 85% of the genes present in any Bilaterian examined were already present in the ur-bilaterian [21], and only about 15% are of more recent origin. Deep comparative genomics including non-bilaterian Metazoans, also revealed that the major developmental signaling pathways are already present in Cnidarians, Ctenophores, Placozoa and sponges [22, 23, 24], and must predate the origin of Bilateria. At the same time, it was shown that genes specific to large clades with divergent sub-taxa, like the deuterostomes, might exist [25]. We hypothesised that similar to the conserved developmental transcriptome and the gene novelties described to define the deuterostomes [25] there might be genes defining the Bilateria as a clade, setting them apart from their non-bilaterian ancestors.

Using reciprocal blast searches, orthology pipelines, and an array of filtering steps we identified a set of orthologues proteins that satisfy our initial criteria for Bilateria specificity. However, these proteins could not be determined to be actually clade-specific for Bilateria with highest-confidence after secondary validation. In fact, re-analysing an initial set of 85 orthogroups with new data from the ever growing database of available proteins we found only 35 orthogroups to remain “Bilateria-specific”. It must be assumed that this set would shrink even further with more data becoming available, in particular from more non-bilaterians.

Despite our limited confidence in the ability of these approaches to detect clade-specific genes in Bilateria, we observed an interesting signal in our data. Analysing the set of 85 orthogroups in more detail we found that these genes are not only retained in species separated by a billion years of independent lineage evolution, but that most of them act in key developmental processes. These genes show very similar and peculiar expression profiles across analogous developmental stages in highly derived model species in the fishes, the arthropods, and the nematodes. This finding supports the idea that evolutionary changes in the genetic machinery of development were coupled to the emergence of Bilateria and are still conserved in very divergent taxa today. Additionally, by finding similar developmental signals across the expression profiles for the most conserved orthologues in our orthogroups, our work also lends new support to the ortholog conjecture.

## 2. Results

### 2.1. Defining potential clade-specific proteins in Bilateria

The aim of our study was to identify genes that newly emerged in the ‘ur-bilaterian’ (Fig 1a) and have been retained in its descendant species.

**Figure 1:**
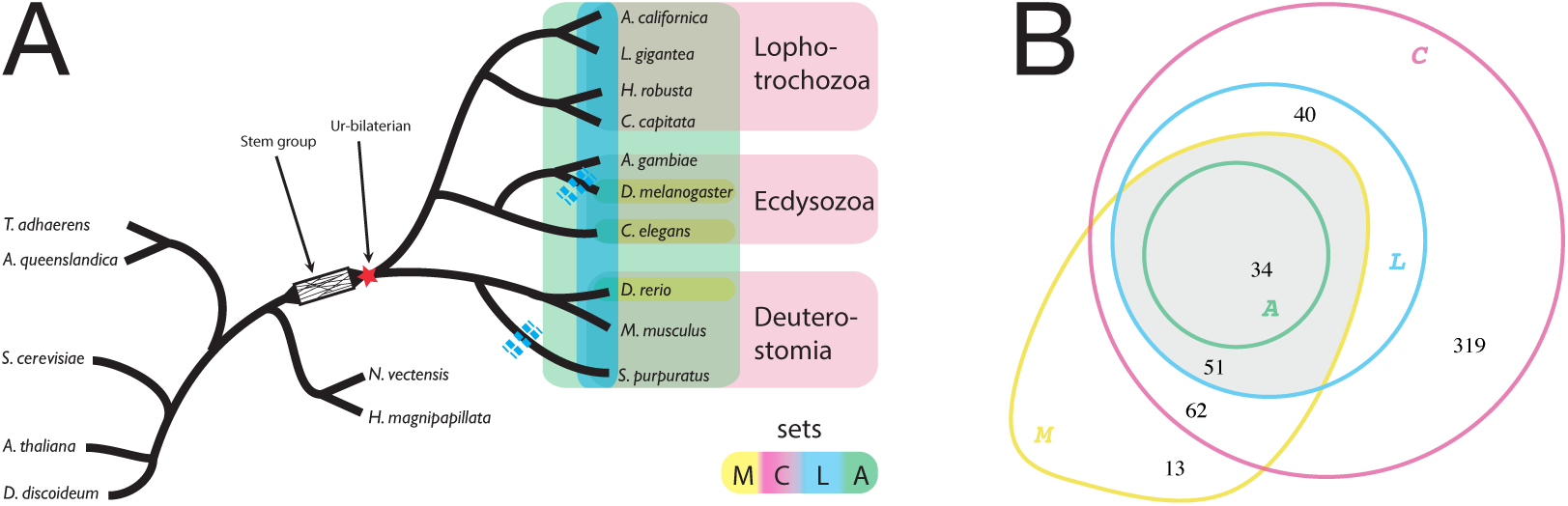
A: Sketch of the genealogical relationship between the clades of Lophotrochozoa, Ecdysozoa and Deuterostomia and the species included in our analysis (Table 1). Legend: *M* – all <u>m</u>odel species are represented in a cluster; *C* – all <u>c</u>lades are represented; *L* – at most one <u>l</u>oss event along the tree (see supplementary Table S1); *A* – <u>a</u>ll of nine bilaterian species are represented. Sketch of the ’ur-bilaterion’ after [4]. Blue dashed lines indicate examples of gene loss events. B: Absolute numbers of clusters of orthologous proteins (COPs) in the four sets. Gray region: detailed analyses are performed for 85 COPs in set *L*′ = *L* ∩ *M*.

**Figure 2:**
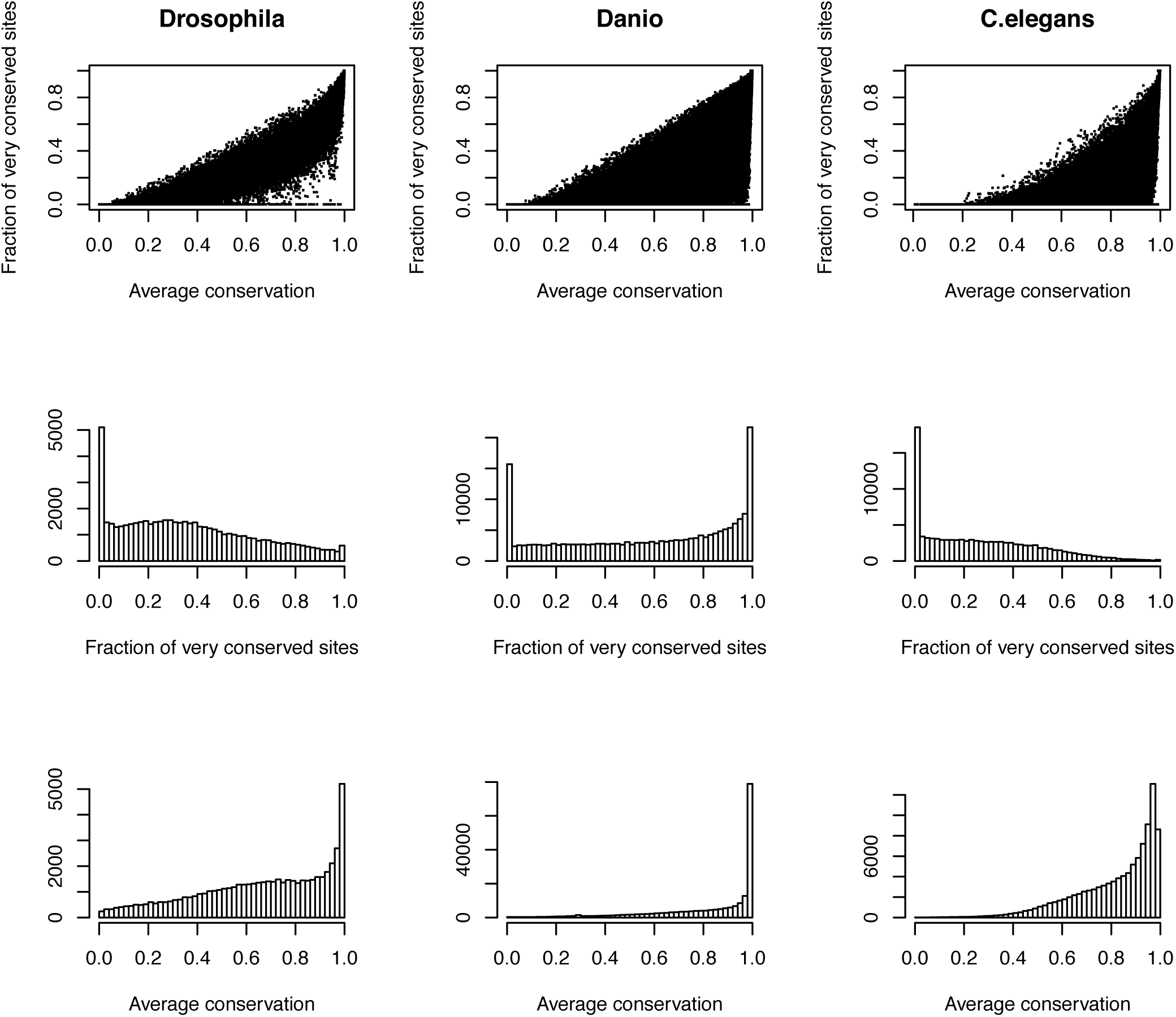
Conservation scores for ortholgous exons from PhastCons analysis. For each species, we show the correlation between scores (top), the distribution of the fraction of very conserved sites (middle), and of the average conservation score (bottom) among exons.

**Table 1:**
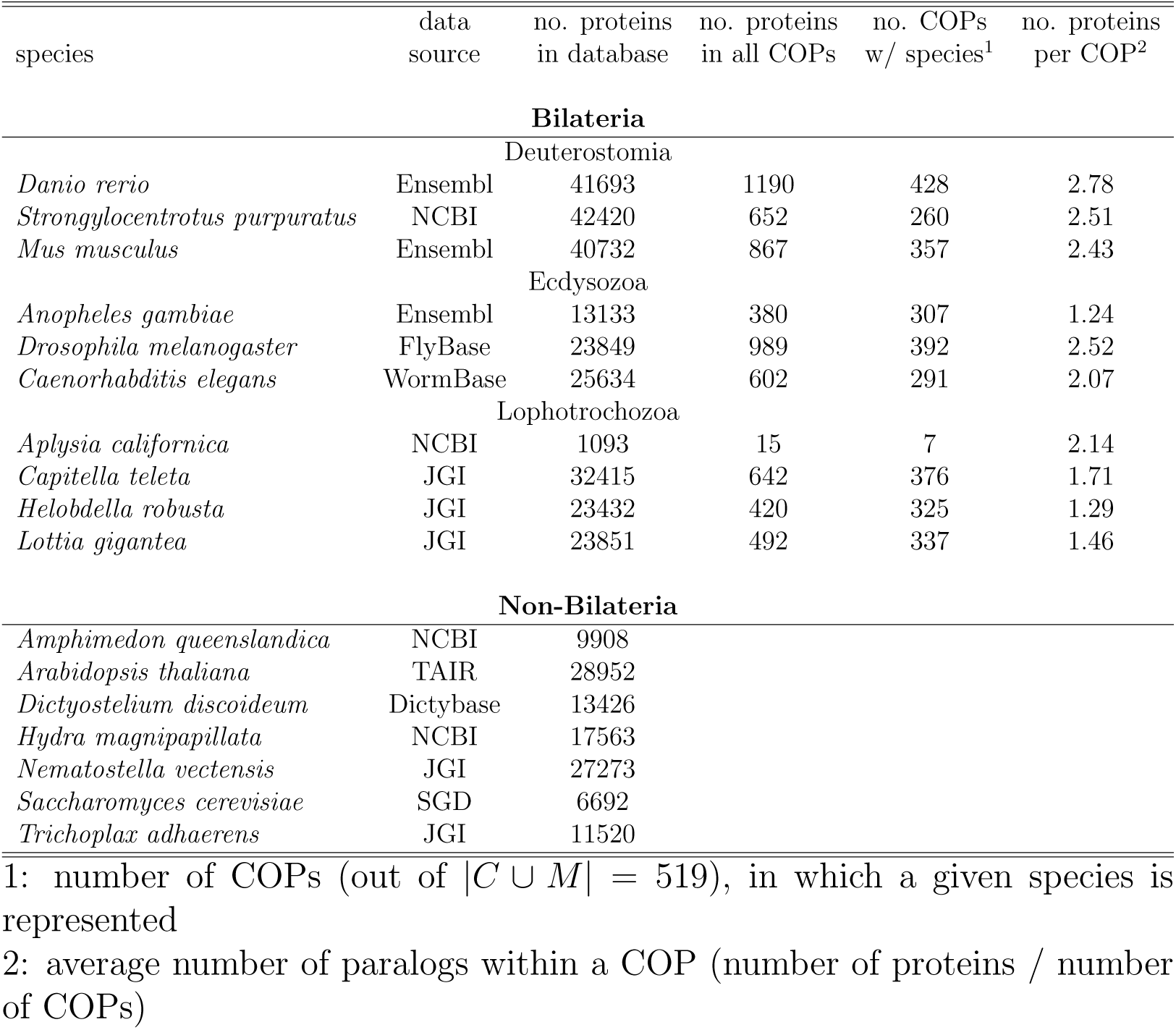
Species considered to contrast bilaterian and non-bilaterian protein repertoires

To this end, our initial study was grounded in a comparison of ten bilaterian and seven non-bilaterian species with fully sequenced and annotated genomes as of 2011. We downloaded 383, 586 protein sequences in total. About 70% (268, 252) are from Bilateria, covering the three major clades Lophotrochozoa, Ecdysozoa and Deuterostomia, and about 30% (115, 334) are non-bilaterian (Table 1). We discarded all bilaterian sequences that had a best BLAST hit below the threshold *E*-value of 10*^-^*^5^ with non-bilaterian sequences. We retained 13, 582 bilaterian specific candidate proteins which were grouped into 1, 867 clusters of orthologous proteins (COPs) using the OrthoMCL pipeline [26]. We condensed the raw set of 1, 867 clusters of orthologous proteins by applying filters with different stringency for the taxa included (Fig 1) and obtained the following four sets

*C*: Each cluster must contain one or more representatives from each of the three major **c**lades, Lophotrochozoa, Ecdysozoa and Deuterostomia. This set contains 506 clusters.

*M*: Each cluster must contain representatives from all **m**odel organisms, *D. rerio*, *D. melanogaster* and *C. elegans*. This set contains 160 clusters.

*L*: Each cluster contains representatives from all major clades, as in set *C*. However, only those clusters are permitted for which the representation of species is explained by at most one **l**oss event along the species tree (set *C* does not have this restriction). This resulted in 125 clusters (see supplementary Table S1)

*A*: Each cluster contains representatives of **a**ll bilaterian species considered (no protein loss is admitted) except possibly *A. californica*. This set has 34 clusters.

On average, each of the nine bilaterian species is represented in 66.2% of the 506 ortholog clusters in set *C*. In set *M*, this percentage is 80.9%, and in set *L* it is 91.2% (see Table 2), reflecting the levels of stringency of the filtering criteria.

**Table 2:**
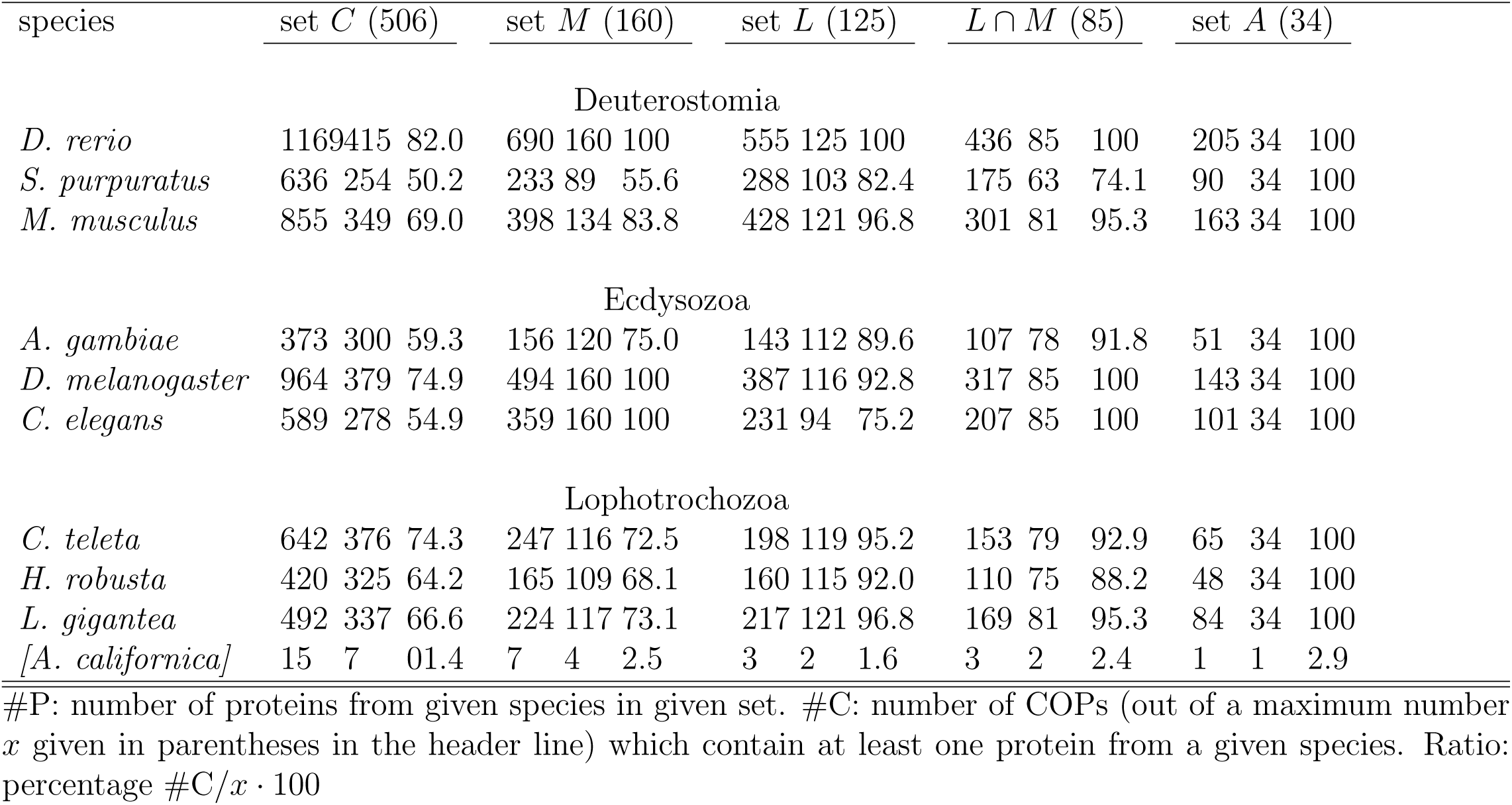
COP distribution in different data sets

### 2.2. Orthologues conserved across divergent Bilateria

In our analysis, we included the highly divergent bilaterian model organisms *Caenorhabditis elegans*, *Drosophila melanogaster*, and *Danio rerio*.

These have very well curated and annotated genomes and are therefore helpful to find genes which are also functionally conserved since more than 500Myrs of independent evolution. To take advantage of this, we decided to use the intersection of sets *M* and *L* (termed *L*′ = *L ∩ M*) for further analysis. Compared to set *L* (125 clusters), this intersection lacks 31 clusters missing a *C. elegans* ortholog and 9 clusters missing an ortholog from *D. melanogaster*, resulting in a set of 85 orthogroups. Except for *A. californica*, the least represented species in these clusters is *S. purpuratus* (genes of this species occur in 63 of 85 clusters). By construction the model organisms (*C. elegans*, *D. melanogaster*, *D. rerio*) are represented in all clusters in *L*′. On average, a (bilaterian) species is represented in 93.1% of the clusters.

### 2.3. Clade-specificity declines with data availability

A general drawback of our experimental procedure is its reliance on correctly identified and annotated genes. Since the majority of the organisms in our study are non-model organisms, they may suffer from incomplete or erroneous gene annotation. For example, we omitted the bilaterian species *A. californica* from downstream analyses because of its low quality protein annotation. It is also possible that truly existing genes are missing from the non-bilaterian dataset due to annotation or prediction errors or simple insufficient availability of sequenced species, leading to wrong inferences of bilaterian specificity. To test this possibility, we first verified by additional BLAST searches at NCBI that none of the 85 COPs in set *L*′ had BLAST hits below an *E*-value of 10*^-^*^5^ in non-bilaterians or other eukaryotes available in August 2014. To accomodate to the very fast growing amount of data from across the bilateria and non-bilateria becoming available in recent years, we decided to implement a second, more comprehensive test to validate the potential bilaterian specificity set *L*′. Conducting searches with Diamond and NCBI-BLAST against a wide array of protein data downloaded in August 2017 from NCBI we identified potential homologs for proteins in the 85 COPs. Using OMA and phylogenetic inferences we could then identify non-bilaterian orthologs for all but 35 of the 85 set *L*′ COPs. This showed that with additional genomes and transcriptomes becoming available, the number of bilaterian specific orthologous groups decreased drastically.

#### 2.3.1. Most conserved orthologues

Althoug not being strictly bilaterian-specific, we regarded the set *L*′ COPs as biologically interesting since they are conserved across huge evolutionary distances with many speciation events separating them in the three model organisms (and additionally divergent enough from non-bilaterian sequences to not be detected in our first re-BLAST screen). We thus wondered what their biological function is and if this function could be correlated to a distinct process in the biology of Bilateria. As a result of our experimental design, but also due to different duplication histories, all COPs contain differing numbers of paralogs from different species. On average, we found most paralogs in Deuterostomia, followed by Ecdysozoa and Lophotrochozoa. The highest average paralog number is found for *D. rerio*, which might be a consequence of the additional genome duplication events in Teleosts [27]. Since the function of paralogs might differ from the function in the last common ancestor, we constructed an additional set of ortholog clusters free of paralogs: we extracted from each cluster in our original set *L*′ the most conserved or-tholog for each species, according to the PhastCons conservation score [28], and discarded the less conserved paralogs. This reduction does not affect the number of clusters, but the number of proteins within a cluster. In our down-stream analyses we considered both versions, the ‘most conserved orthologs’ (MCO) and the ‘all orthologs’ (AO) catalog for each cluster.

### 2.4. Gene ontology and functional classification GO terms reveal a link to development

To gain an overview over the biological functions of the proteins in set *L*′, we extracted and analyzed their associated GO terms. To quantify simultaneous enrichment in multiple species, we developed a novel generalized Fisher’s exact test for multiple species, which we term MSGEA (‘Multi Species GO Enrichment Analysis’; Fig 3 and Fig S1). In contrast to standard GO enrichment analysis, MSGEA does not solely focus on the over-representation of GO terms in single species, but is able to detect GO terms enriched across several species, even if not over-represented in any single species. Of special interest for our analysis were GO terms occurring across all three model organisms: such terms may indicate the conservation of biological function across large evolutionary time scales. To our knowledge, this novel method is the only test that is sensitive to coherent enrichment of GO terms across multiple species (see Methods). Applying MSGEA we extracted a list of GO terms which are likely associated to long-term retained functions of our candidate genes. Concentrating on the domain ’biological process’ of the GO database, we find the terms ‘development’, ‘muscle’, ‘neuron’, ‘signaling’ and ‘regulation’ to be strongly over-represented (Fig. 4 and supplementary Table S4). The terms identified through in silico analysis suggest a prominent role for these orthologues in development.

**Figure 3:**
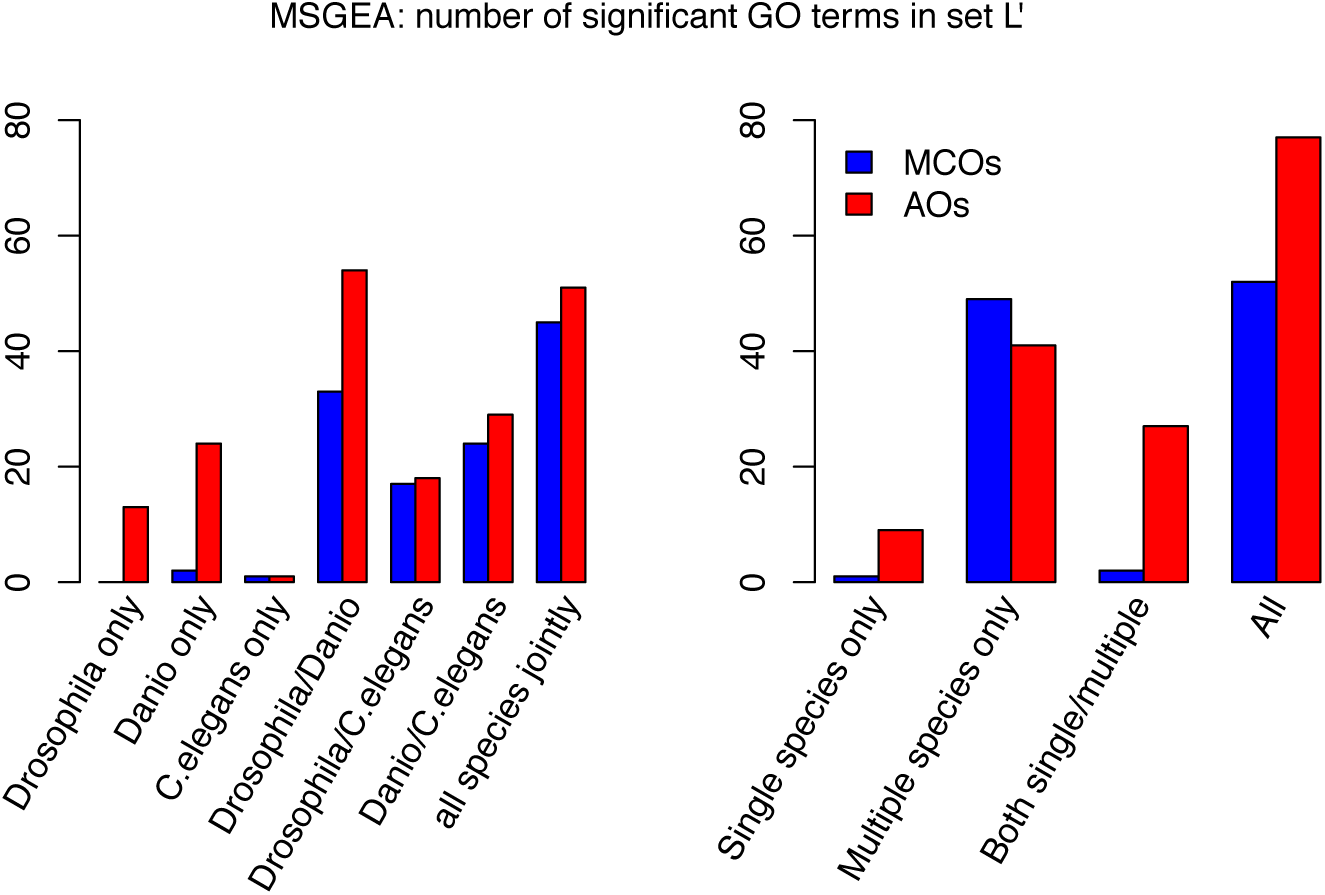
Number of significant GO terms identified with MSGEA in set *L*′ (*p <* 0.05). Red histograms: considering all orthologs (AO) from a cluster; blue histograms: only the most conserved orthologs (MCO) are considered in the MSGEA analysis. Left: number of significant terms found for each single- or multi-species enrichment analysis. Right: number of significant terms found only in single/multiple-species enrichment analyses and in both kind of analyses. The large amount of GO terms found only in multiple species analysis illustrates the power of MSGEA (see also Fig S1).

**Figure 4:**
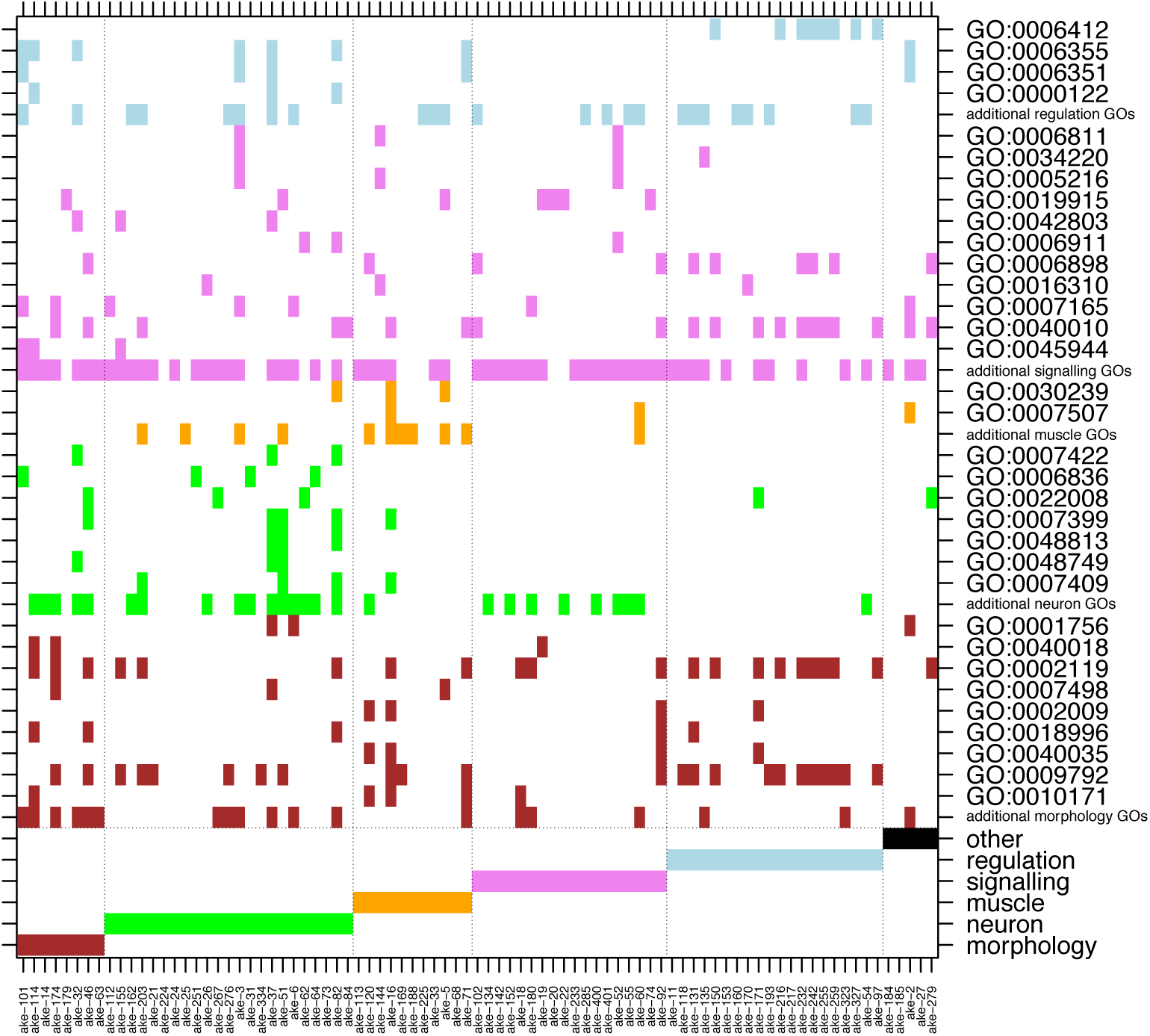
Significant GO terms identified in set *L*′ by applying MSGEA (see text). The upper part of the figure shows the GO terms which occur in each COP (*x*-axis). Those terms which occur at least in three COPs are spelled out with their GO-ID, otherwise they are collected in the lines ‘additional … GOs’. Only non-generic GO terms of level 4 and higher are recorded. The bottom six rows show the assignment of COPs to functional classes based on human-curated literature mining. As can be seen from the color code, automatic GO and human-curated functional assignments do not always coincide.

#### Inference of biological function through literature mining

Since orthology is a strictly phylogenetic criterion, and since GO classification draws heavily on sequence homology, without necessarily reflecting conserved function, we decided to perform an in-depth human-curated literature search to collect additional evidence for the functional role of genes in COP clusters. We mined the literature databases and compiled human curated functional descriptions, based on experimental evidence, for the proteins contained in set *L*′. We defined the six classes ‘Neuron related’, ‘Morphology related’, ‘Muscle related’, ‘Signaling related’, ‘Regulation related’ and ‘Others’ (supplementary Table S5). We used functional descriptions extracted from the literature to assign each of the 85 set *L*′ clusters to one of these six classes (Fig. 4). The manual extraction of species-and protein-specific functional information allowed us to compare and better interpret functions across the three model organisms.

To compare this human-curated classification with the one based on the GO database, we assigned all GO terms appearing in set *L*′ to one of the six classes above. Since a given protein may be associated with many GO terms there is no one-to-one relationship of the human curated annotation and the GO terms. For example, while the human curation assigns eight of the 85 clusters (the left-most columns in Fig 4) to the category ‘Morphology related’, morphology related GO terms occur in 33 clusters, scattered throughout the six classes. Similarly, signaling related GO terms are found in almost every cluster, while the class ‘Signaling related’ contains only 18 clusters. However, our manual curation is still broadly consistent with GO annotations. For example, GO terms associated to ‘muscle’ do occur in the class ‘Muscle related’ and GO terms related to ‘neuron’ are mostly found in the ‘Neuron related’ clusters.

### 2.5. Expression profiles

To analyze expression of genes in set *L*′ across developmental stages and to compare it between model organisms, we used publicly available data for *D. melanogaster*, *D. rerio* and *C. elegans*. We normalized expression values - separately for each species and for each developmental stage - by the genome-wide average, and then log-transformed them.

#### Pooled profiles

To obtain a global picture of the expression profiles of genes in set *L*′, we calculated mean and median profiles for each of the three model organisms, shown as orange lines in Fig 5 and Fig S3, top row. To see how expression of bilaterian-specific genes compares to background expression, we generated two different background sets for each species. The first is a random-background, obtained by randomly selecting genes from the model organism genomes and recording their expression (distribution shown as white box-plots in Fig 5). The second background, which we call the ‘matched-function’ background (orange boxplots), is a random collection of genes with GO terms matched to those found in set *L*′. The background distributions were constructed independently for each stage and for each set, and they may differ in size and content. We grouped the developmental stages from fertilized egg to the adult body into coarse grained, yet comparable, phases: ‘Blastoderm’, ‘Gastrulation and Organogenesis’, ‘Hatching to Larva’ and ‘Adult’ (separated by vertical lines in Fig 5).

**Figure 5:**
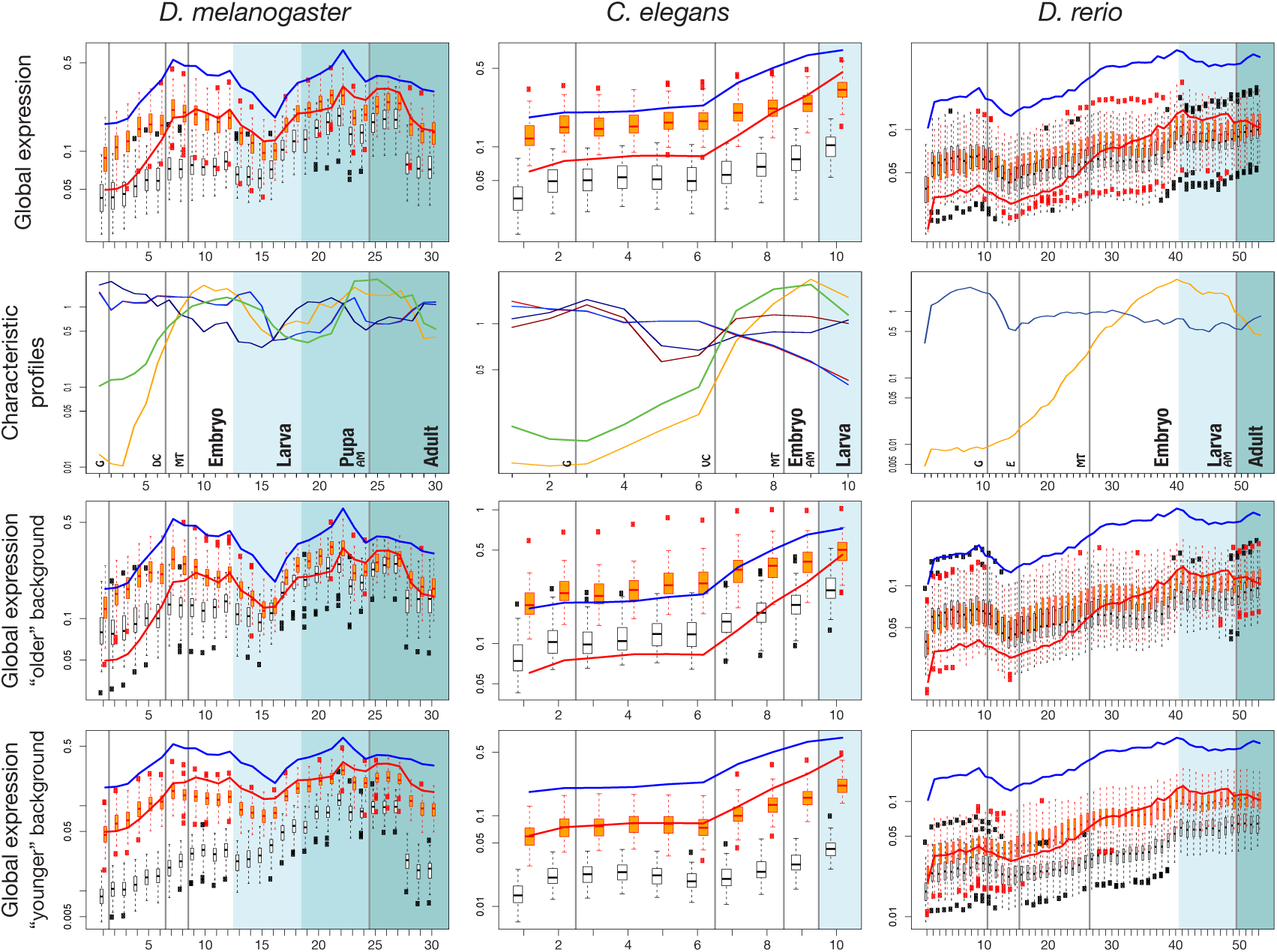
**Expression profiles** of different species. Top row: average (global) expression profiles for genes collected in set *L*′. Red line: pooled profiles for ’all orthologs and paralogs’ (set ’AO’). Blue line: profiles for the ’most conserved orthologs’ (set ’MCO’). The white and orange boxplots show the ’random’ and ’matched-function’ expression backgrounds, respectively (see text). Second row: Characteristic expression profiles. Different colors represent different clusters. Darker and lighter colors represent profiles which are similar among species. Bottom rows: distributions shown in boxplots are calculated from genes which are younger (phylostrata ≥ 7 according to the terminology of [32]) or older (phylostrata *<* 7) than bilateria. Background shading (from white to dark grey) represents developmental stages embryo, larva, pupa (fly only) and adult. For a detailed explanation of the developmental stages see supplementary Tables S2 and S3. Meaning of labels in row 2: G - gastrulation; DC - dorsal closure; MT - first muscle twitching; AM - transition to adult morphology; VC - ventral closure; E - 100% epiboly

For all three species, the set *L*′ profiles roughly follow the shape of the background profiles across embryogenesis, however at a higher expression level (Fig 5). In *D. melanogaster*, expression of bilaterian-specific proteins is initially low, then rises until ‘dorsal closure’, after which it slighly decreases again. In *C. elegans*, expression is also initially low and then rises towards ‘ventral enclosure’. This pattern is mirrored in *D. rerio*, where expression peaks around the pharyngula stage. The profiles seen in *C. elegans* and *D. rerio* are congruent with the major cycle of tissue proliferation and differentiation during organogenesis in these species [29]. During ‘dorsal closure’ in *Drosophila*, ‘ventral closure’ in *C. elegans* and ‘pharyngula’ in *Danio*, the head, the nervous system and the bilaterally organized body develop [30]. In *Drosophila* we find a second expression peak during a second cycle of proliferation corresponding to metamorphosis. The matched-function profile largely resembles the profile of set *L*′. However, set *L*′ genes show distinctly low expression in the very early embryonic stages, a pattern which is consistently seen in all species (Fig 5 and Fig S3). Another common feature is that expression of MCO genes is consistently higher than of AO genes (Fig 5).

#### Characteristic expression profiles

To further analyze the expression profiles, we clustered individual profiles based on their similarity and extracted a few typical ones for each species (Fig 5, bottom row). Methods exist to identify tissue-specific expression profiles, or to differentiate between profiles, but no standard techniques are available to detect enrichment of profiles across a set of genes. To overcome this limitation, we employed an ad hoc ’profile enrichment’ strategy. Briefly, we considered a COP expression profile relevant, if similar profiles were significantly more abundant among genes in our COPs than among other genes (for details, see Methods). We extracted all statistically significant profiles and then clustered them to distill representative ones. For each of the three model species qualitatively similar patterns are retrieved: an intersection of increasing expression profiles (yellow and green lines in Fig 5, 2^nd^ row) with slightly decreasing profiles (dark red and dark blue lines). The intersection occurs right before the developmentally important events ‘dorsal closure’, ‘ventral enclosure’ and ‘pharyngula’. A second crossing of profiles can be seen in the fly, just before metamorphosis into the adult body. To interpret this observation, we considered the following functional classification.

#### Expression profiles stratified by function and age

We determined the mean expression profiles for the six functional classes - morphology- (8 set *L*′ COPs), muscle- (11), neuron- (23), signaling- (18), regulation-related (20) and other genes (5) (Fig S2). We find upward and downward peaked expression patterns for the classes morphogenesis, neuron and muscle development (Fig S4), which are even emphasized compared to the pooled profiles of the global analysis. This is in particular true for expression before the onset of gastrulation. On the contrary, the genes related to regulation and signaling display a rather constant profile, without the characteristic expression peaks of morphogenesis.

For species as different as animals, plants and fungi, it has been repeatedly observed that timing and level of expression during development depend on the evolutionary age of a protein [31, 32, 33, 30, 34]. For instance, one recurrent finding was that older genes tend to show higher expression during early embryogenesis. To examine our proteins in this respect, we analysed whether the set *L*′ expression profiles are more similar to the profiles of younger or of older genes, and we calculated background distributions considering the age of genes. Genes which arose before Bilateria (phylostratigraphic level less than 7, corresponding to Bilateria, according to the strata-numbers by Domazet-Lošo) were defined as ’older’ and all others (level *>* 7) as ’younger’ [32]. Expression profiles of genes (AO) in set *L*′ match very well the profiles of the ’younger’ genes with matched function (red lines and orange boxplots in Fig 5 and Fig S3). All three species show low expression of young and of set *L*′ genes during early development. Hence, this late onset of expression appear to be an evolutionary novelty of Bilateria. After ‘dorsal closure’ in *D. melanogaster*, ‘ventral closure’ in *C. elegans*, and ‘pharyngula’ in *D. rerio*, expression increases and older and younger profiles converge.

## 3. Discussion

The most significant innovations of bilaterian animals are a third germ layer (the mesoderm), a liquid filled body cavity surrounded by mesoderm (the coelom), a complex nervous system, and a large number of specialised cell types. The formation and presence of these features had a massive effect on the evolutionary possibilities and success of bilaterians. By analysing the genomes of bilaterian and non-bilaterian species we aimed at testing whether there is a genomic correlate to these morphological key innovations. Roughly a billion years of divergence between the most distant species, chromosome fusions, fissions and re-arrangements, genome duplications, expansions and losses of gene families, as well as myriads of nucleotide substitutions might have blurred the common genetic heritage of Bilateria. However, those genes and proteins which have evolved under steady purifying selection, i.e. play a key non-redundant role in extant Bilateria species, should still be identifiable as orthologs across Bilateria. We searched for genes that potentially evolved in the bilaterian ancestor and have been retained in extant bilaterians because such genes might have influenced the evolution of bilaterians. We were initially able to detect a set of 85 orthogroups (set *L*′), containing genes which are shared between widely divergent Bilateria and, at the same time, appeared to be without orthologues in non-Bilateria. We reason that this set had to be taken with caution, as for example there is no clear-cut threshold of sequence similarity which separates orthology from non-orthology. In such cases the decision, whether a gene with partial homology should be included into an orthogroup or not, is somewhat arbitrary. For the 85 set *L*′ COPs we have considered partial homology to a non-bilaterian sequence as an insufficient criterion to exclude an orthogroup from our initial list. A similar BLAST strategy as implemented by us, however without subsequent orthology clustering, had been used before to determine the evolutionary origin of genes and to assign phylostratigraphic ages [35, 32, 36, 34]. One pitfall of this approach is that an overall alignment score, such as an Blast *E*-value, may lead to both false positive and false negative predictions. False positives arise from the (wrong) inclusion of a protein into the bilaterian-specific list, which *does* have non-bilaterian orthologs, but which are missed due to low sequence similarity or due to absence from the database. False negative predictions arise from missing truly bilaterian-specific proteins, for instance when a conserved domain is identified in a protein for which a homologous domain is also present in non-bilaterians. One such example is the C2H2 zinc finger protein CTCF which has been described as bilaterian-specific before [37]. Our search did not recover this protein, because multi-zinc finger proteins exist in cnidarians and their conserved Cys/His residues and linker regions evoke a similarity above threshold. As a result, our approach will wrongly treat such proteins as older and not bilaterian-specific and, consequently, eliminate them. Similar errors may affect other proteins with extended conserved domains. However, proteins that pass the filter should have a high probability to be true bilaterian novelties. In contrast, genes may be wrongly assigned to subclades which are younger than bilateria. For instance, this could be a consequence of non-orthologous gene displacement (NOGD) [38], a process in which functions are taken over by new or different proteins and the original orthologue is lost. Finally, we could have wrongly classified a protein as bilaterian-specific because faint homology to more anciently diverged species is missed.

Another limitation in our initial screen was the restriction to 16 high-quality genomes, from which we infered bilaterian specificity. Firstly, it has now been shown that OrthoMLC, which we mainly relied on, is limited in finding distantly related orthologues if taxon sampling is coarse [39]. This problem could have been inflated by our restriction of candidates in our blast searches before running orthology prediction pipelines. Secondly, new entries to the databases may change the inferred phylostratigraphic age of an orthogroup. Being aware of this and in particular because sequencing efforts are increasingly directed towards non-standard model organims in the last years, we have scrutinized and re-inspected our initial candidate set of set *L*′ COPs with refined tools and new data. While the remaining sub-set of 35 orthogroups might consist of bilaterian-specific proteins, it is likely that more data from so far not-yet-sequenced non-bilaterian species will also reveal orthologous for some of these proteins. These caveats raise doubts on the likelihood to detect high-confidence genes that are unequivocally bilaterian and at the same time widely retained in this taxon. Our results clearly raise the possibility that there are just a handful of Bilateria-specific genes, or even none at all. At the same time our approach of partially falsifying the results of the first orthology screen by implementing two additional checks shows that if there is any such gene, careful and elaborate measures have to be taken to validate potential candidates. At the very least all available genomic and proteomic data from currently available databases have to be screened.

Despite not being necessarily clade-specific, our set of genes is retained in very distantly related bilaterian species, while at the same time not being restricted to the group of classical, universally-conserved house keeping genes. This is interesting to note since even highly conserved and important genes can become lost, such as some Hox genes, e.g. from *C. elegans* [40] and the nematodes in general. Our set is also significantly divergent from potential non-bilaterian orthologs, suggesting either their origin or their rapid evolution in the ur-bilaterian. It thus appeared logically to ask which functions these orthologues might have in bilaterian animals. In comparison to classical enrichment analysis implemented in various software packages like blast2GO [41], our MSGEA test allows for the first time to compare GO-enrichment across several species. This is advantageous in the age of genomics where very often more than one species or a number of samples is analysed. In our framework of three model bilaterians, a fruit fly, a zebrafish, and a roundworm, the MSGEA based GO term analysis and subsequent literature mining indicated roles in development for the clustered genes. This link appears likely, since the process of development, the construction of adult specimens from single cells, is conserved in all animals and is also the phase when the combined hallmarks of bilaterian life are brought together: bilateral symmetry, triploblasty, the enhanced nervous system, and a set of complex cell types. In particular, the ‘muscle’ and ‘neuron’ class of clusters is compatible with the idea that key bilaterian innovations involved the central nervous system and muscles [42]. In general the connection to development is in agreement with other studies, such as the large scale modEncode study of model organisms [29].

The conserved genes in set *L*′ follow characteristic expression patterns during development, which are shared between species separated by a billion years of independent evolution. In particular, we observe characteristic low expression in early development and a peak towards the end of larval development common to highly derived taxa such as nematodes, insects and teleosts. The single (fish and nematode) and double peak (fly) patterns which we observed are in agreement with results from the modEncode project [43, 29]. Starting from co-expression analyses, the authors arrived at a set of orthologous genes acting in comparable stages of *D. melanogaster* and *C. elegans* development. The simpler, single peaked pattern in *C. elegans* and *D. rerio* reflects up-regulation at ventral enclosure and pharyngula stages, respectively. Thus, the expression profiles in these organisms mirror their life-cycle, displaying shared peaks when the adult body-plan is assembled. This phase, dominated by morphogenesis, has been interpreted as the phylotypic stage in both organisms [44, 45]. In the holometabolous fly the situation is more complex: a body morphology is built twice, for the larva and then for the adult. These are two phases of increased cell proliferation during gastrulation and the transition from pupa to adult. The two peaks are also visible in the matched-function set, but they are less pronounced in the random-background set. It thus appears the two peaks we observed for our proteins reflect up-regulation during both phases of cell proliferation. It will be interesting to analyse if similar patterns can be observed in other species undergoing metamorphosis, particularly in vertebrates.

It has been hypothesised that differences in gene regulatory processes and the deployment of GRNs may have been more important for the evolution of Bilateria than the conservation of single key genes [46]. A conserved profile of developmental gene expression was observed in several studies across animal phyla, including in non-bilaterians, when developmental time course data were analysed [12]. These studies suggest the existence of a phylotypic stage late in development, but see [47, 12, 31]. We observed low expression of young genes that originated after the emergence of Bilateria and of set *L*′ genes in early development. We interpret this to indicate that early embryonic processes, e.g. initial cell divisions, are governed by evolutionary old genes, which are common to all bilateria and non-bilateria. Thus while, non-Bilateria have a similar complex distribution and abundance of gene regulatory elements and systems [48, 49], and likely their own phylotypic stage, our data support the idea that a change in gene regulation in the phase when adult morphology is shaped might be linked to the emergence of Bilateria. What is more, the process appears to be conserved across immense evolutionary distances.

In agreement with the general idea of the orthology-conjecture, paralogs contained in set AO may be sub-functionalized, while the MCO genes are more likely to retain their original function. This observation on the MCO genes appears interesting in regard to the ortholog conjecture, which posits that orthologues are more likely to retain function in divergent organisms in comparison to paralogues, see e.g. [50]. While being challenged in the past [51] more recent analyses have found support or at least some support for the ortholog conjecture by comparing for example mammals or divergent model organisms [52, 53, 54]. Our data does give additional support to the idea of functional conservation of orthologues in very distant species and in particular potentially extends the conjecture to expression patterns. That paralogues are indeed more divergent in their function was for example observed in flies [55] and appear apparent in our data of the AO subset as well.

### 3.1. Conclusions

In this study we performed a systematic approach to identify bilaterian-specific genes. Previous investigations concluded that essentially all important developmental regulators precede the origin of bilaterians, and that rewiring of already existing factors was a hallmark of bilaterian evolution [24, 56, 23, 22, 48] and our results enhanced by a second BLAST and phylogeny based control step are in support of this. Future data from non-bilaterian species might in fact show that even the 35 groups of orthologues we now identified to be conserved across the three model species but without credible non-bilaterian orthologues were present before the split from Cnidaria. Conversely, it remains possible that genes independently lost in bilaterian crown groups have no non-bilaterian orthologues. Analysing the roles of genes we found conserved across crown groups in bilateria we found a strong link to development. Thus, our results indicate that Bilateria are unified by processes occuring late in embryonic development, which are shaped by a set of conserved genes. At the same time early developmental processes might be shared across Metazoa. Finally, our findings give new support to the ortholog conjuncture.

## Materials and Methods

Using a comparative genomic approach, we aimed to identify genes shared among and specific to Bilateria. To represent the spectrum of the different bilaterian clades we selected ten species from the three major clades - Deuterostomia, Ecdysozoa and Lophotrochozoa - concentrating on species with a fully sequenced and annotated genome (Table 1). We included more species from Lophotrochozoa than from the other two clades, since well-annotated complete genome sequences from lophotrochozoan model-organisms were still under-represented in public databases at the beginning of our studies. To obtain a high contrast between Bilateria and Non-bilateria, we used seven non-bilaterian species, as in a previous study [37]. Proteomes and data from InterProScan [57] and the Gene Ontology consortium [58] were retrieved from the sources cited in Table 1.

### Initial BLAST and Clustering

To find orthologs we set up a stringent analysis pipeline. We first species blasted all-versus-all among the 17 proteomes using the stand-alone BLAST version 2.2.25+ with a cutoff of *E* = 10*^-^*^5^. We then discarded all bilaterian proteins which had a significant hit in any of the non-bilaterian sequences.

This filtering resulted in 13, 582 candidate proteins. From these we constructed orthologous groups using two different ortholog finders, InParanoid and OrthoMCL [26, 59], using default parameter settings. Both programs were rated highly in benchmarking studies which analyzed the performance of orthology-prediction methods [60, 61]. For the purpose of detecting orthologs of very diverged species, we chose OrthoMCL version 2.0.2 and the associated MCL version 11-335, as they were shown to be more robust [60, 61] than similar programs. Finally, we grouped the ortholog clusters into four sets (*A*, *L*, *M*, *C*; Fig 1), satisfying different stringency conditions, see page 4. Due to the poor sequence quality, we treated the Lophotrochozoon *A. californica* differently: where available we included *A. californica* orthologs, but we did not require them to be present in set *A* (’all species’).

The above data sets are ordered by degree of initial confidence in the bilaterian specificity of the respective proteins. At the same time they reflect the level of conservation of the proteins across different species. Seeking to be conservative regarding bilaterian specificity on the one hand (set *A* being most conservative), but on the other hand to analyze a dataset which is as comprehensive as possible, we focused our further analyses on the intersection of sets *L* and *M* (called *L*′; Fig 1) comprising 85 clusters.

### Validation with BLAST, OMA, and Phylogeny

As our original dataset was bilaterian and non-bilaterian species was assembled prior to 2012 and the steady flow of newly sequenced species has added massive new numbers of protein predictions to online databases we wanted to re-evaluate the potential bilaterian-specificity of our 85 final clusters. To this end we downloaded 19,185,382 bilaterian and 497,273 non-bilaterian proteins from the NCBI database in August 2017 and created individual databases for taxa in bilateria and non-bilateria. Next we used Diamond [62] to blast the *D. rerio*, *C. elegans*, and *D. melanogaster* proteins for each of the 85 clusters against the individual taxa. Additionally, we used NCBI BLAST with the parameters described to be most sensitive in finding orthologs in [63] for the non-bilaterian taxa. For each of the 85 clusters we then collected the 10 best blastp hits per taxon. Along with the original model organism proteins we obtained 96,287 proteins in this way, which we used as input for the OMA orthology prediction pipeline [64]. From the OMA output we selected hierachical cluster of orthologues (HOG) for those of the original 85 clusters where the HOG contained non-bilaterian proteins. Finally, we aligned all sequences per HOG with clustal-omega [65] and reconstructed a phylogeny using IQTrees with automated model selection and 1000 fast bootstrap replicates [66]. We then analysed these trees by eye.

### Selecting most conserved orthologs

For the three model species *D. rerio*, *D. melanogaster*, and *C. elegans* we extracted the UCSC tracks of basewise PhastCons conservation scores [28] calculated across insects, teleosts and nematodes, respectively. We used these scores to rank all genes in a cluster according to their fraction of strongly conserved sites, i.e. sites with PhastCons score *>* 0.99. We selected the highest-ranking one from each cluster as the ’Most Conserved Ortholog’ (MCO). These genes likely retained the same function, even in highly diverged species – a conjecture based on the idea that strong conservation reflects long-term evolutionary (and functional) constraint and that neo- or sub-functionalized paralogs tend to be less conserved. We used the fraction of strongly conserved sites, instead of the average conservation score, since we are interested in the degree of conservation in function, not in sequence. We reasoned that functional conservation should be related to high conservation of alleles at functional sites, while the remaining bases can evolve fast. However, the two ways of measuring similarity are highly correlated on a genome-wide scale across all coding exons (*r* = 0.91 for *D. melanogaster*, 0.84 for *D. rerio* and 0.71 for *C. elegans* (Fig 2)). We performed all subsequent analyses on both sets: the set of ’All orthologs and paralogs’ (AOs) and the set of MCOs.

### Multi-Species Gene Enrichment Analysis

To examine function of the genes in set *L*′ by *in silico* methods, we obtained their gene ontology (GO) terms for the three model species from the GO database [58]. To identify terms enriched in COP genes compared to the genomic background, we employed two strategies: the conventional single-species enrichment analysis based on Fisher’s exact test and a novel method, termed multi-species gene enrichment analysis (MSGEA). Standard single-species Fisher’s exact tests for enrichment cannot capture a signal of moderate, but joint, enrichment across different species. Instead, with MSGEA we compute an exact *p*-value for a pooled set of species, employing the following steps. First, for each GO term we counted the number of descendants in the GO tree. Then, we computed the *p*-values for the terms occurring in any species and corrected for multiple testing [67]. More specifically, let *n_go,s_* be the count (i.e., number of occurrences) of the GO term *go* in species *s* contained in our set and let *N_go,s_* be the count for the whole genome. Furthermore, let *n_s_* = Σ*_go_ n_go,s_* and *N_s_* = Σ*_go_ N_go,s_*. Assume that all species under consideration diverged at the same time and evolved independently after that (i.e. they have a starlike phylogeny). The null hypothesis of the MSGEA test is that a given GO term was not over-represented in the common ancestor. Therefore, a significant *p*-value means that the genome of the ancestor was already enriched in the GO term considered. Being conservative, assume that the null distributions of *n_go,s_* are independent hypergeometrics with parameters *N_go,s_*, *n_s_* and *N_s_* in each species. The enrichment statistics *X_go_* is then the sum of the normalized enrichments of *n_go,s_* across species, i.e. the sum of the *z*-scores:

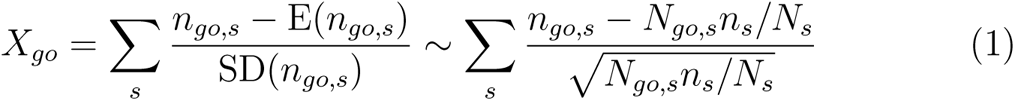

where a Poisson approximation is used to define the score. Finally, the *p*-value is the probability

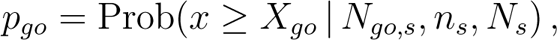

where the distribution of *x* follows from the hypergeometric distributions of the *n_go,s_* with the above parameters.

For a single species, this test coincides with the standard one-tailed Fisher’s exact test for GO enrichment, and therefore is consistent. The exact estimation of *p*-values is computationally intensive; an optimized code, written in C, is available from the authors upon request. Since MSGEA is based on the hypothesis of independent evolution of each lineage, it can be applied only to starlike phylogenies. For the situation considered here, this assumption is met, since the “roundworm-fruit fly-zebra fish” phylogeny is approximately starlike. We find that a considerable fraction (30-70%) of significant GO terms in our data set is detected by MSGEA, but not by single species enrichment.

### Detailed functional analysis of genes from model organisms

To mine literature databases for functional studies in model organisms with a focus on development we employed biomaRt: we used the Bioconductor module [68] biomaRt 2.19.3 to extract information for proteins present in cluster set *L*′. Wormbase release WS220 was queried for *C. elegans* proteins and ENSEMBL 75 for *D. melanogaster* and *D. rerio* proteins. We then searched the literature for experimental validations of protein function. We grouped the results in six major categories, described in Results and Discussion and summarized in supplementary Table S5. These categories represent prominent molecular functions during embryogenesis and development. Based on the retrieved annotations we assigned proteins in set *L*′ to these categories.

### Cross species expression profiles

We retrieved expression data for different developmental stages in *D. melanogaster* [69, 70], *D. rerio* [32] and *C. elegans* [71]. *D. rerio* data for adult stages were not used. The expression profiles were transformed by taking the logarithm to base 10 of the expression levels and then subtracting the log_10_ of the mean expression at each stage.

We devised the following test to find characteristic expression patterns: first, we computed Pearson’s correlation coefficient between expression profiles for different genes. For each profile in each set of the genes from 85 COPs, we performed two types of tests: (i) a Mann-Whitney test on the distribution of correlations, comparing the correlation of the profile with other bilaterian genes versus the correlation with genome-wide profiles, in order to detect profiles that are more correlated with the ones of other bilaterian genes than with the rest of the genome; (ii) for each profile, we classified the remaining genes as “highly correlated” or “not highly correlated” in expression, using correlation thresholds of *r* = 0.5, 0.7 and 0.9; then, we tested for enrichment of correlated profiles among Bilaterian genes by Fisher’s exact test.

A Benjamini-Hochberg correction for multiple testing was applied to the *p*-values resulting from these tests. All the significant profiles (*p<* 0.05) were clustered based on their correlation coefficients *r* by complete-linkage hierarchical clustering implemented in the R statistics software package, using 1 − *r* as distance measure and selecting clusters at height *h* = 0.75. After clustering, profiles were normalized by their average expression across stages, then averaged across each cluster. The resulting profile shapes are shown in row 2 of Fig 5.

### Are expression profiles driven by gene function?

We performed a randomisation test to explore if the mean (logarithmic) expression level of COPs genes could be explained by their function. For each developmental stage we checked if expression correlates with that of randomly selected genes possessing equal or similar GO terms, i.e. we checked if genes of different age, but similar ontology, would show similar expression profiles.

We performed this analysis only on set *L*′.

For this purpose, we randomly sampled from the whole genome 100 sets with the same number of genes as are contained in set *L*′. We did this in two different ways. (i) The first (“random-background”) was a random sampling of genes from the genome. After sampling, we computed the mean normalized expression of the selected genes for each stage. We repeated this procedure 100 times, and thus obtained a distribution of normalized means, represented as boxplots (in white) in Fig 5, Fig S3 and Fig S4. (ii) The second was a “matched-function” sampling: first, for each GO term in the original set, we listed all genes with the same annotation and extracted from this list a number of random genes equal to the number of occurrences of the term; second, we pooled all the resulting lists; third, we extracted from this list a number of random genes equal to the number of genes in the original set. This way, we obtained sets of the same size and function (approximately, at least) as our original set. We applied this to the whole set *L*′ and for all subsets corresponding to the six functional categories described above (Fig S4).

### Age index of proteins

We used the phylostratigraphy from Domazet-Lošo et al. [32] for *D. melanogaster*, *D. rerio* and *C. elegans* to define two age-groups of proteins: (i) older proteins, with an origin in phylostrata 1 to 6; (ii) younger proteins, with an origin in phylostrata 8 and higher. Proteins that we identified as bilaterian-specific (i.e. the ones in our sets *A*, *L*, *M* and *C*) were excluded from the two groups, irrespective of their original phylostratigraphic classification. We repeated all the analyses described in the previous sections, and compared separately our COPs genes with the genome-wide background of older and of younger genes (Fig 5).

## Acknowledgements

We are grateful to Peter Heger (University of Cologne) for his input on previous versions of this manuscript. We are also very thankful to Thomas Wiehe (University of Cologne) for his support in conducting this research, his input on functional profile analysis and many vibrant and fruitful discussions leading to the finished manuscript. We thank Georgios Koutsovoulos (INRA Sophia-Antipolis) and Sujai Kumar (University of Edinburgh) for help with the initial orthology clustering.

Supplementary Figures and Tables

**Figure S1.**
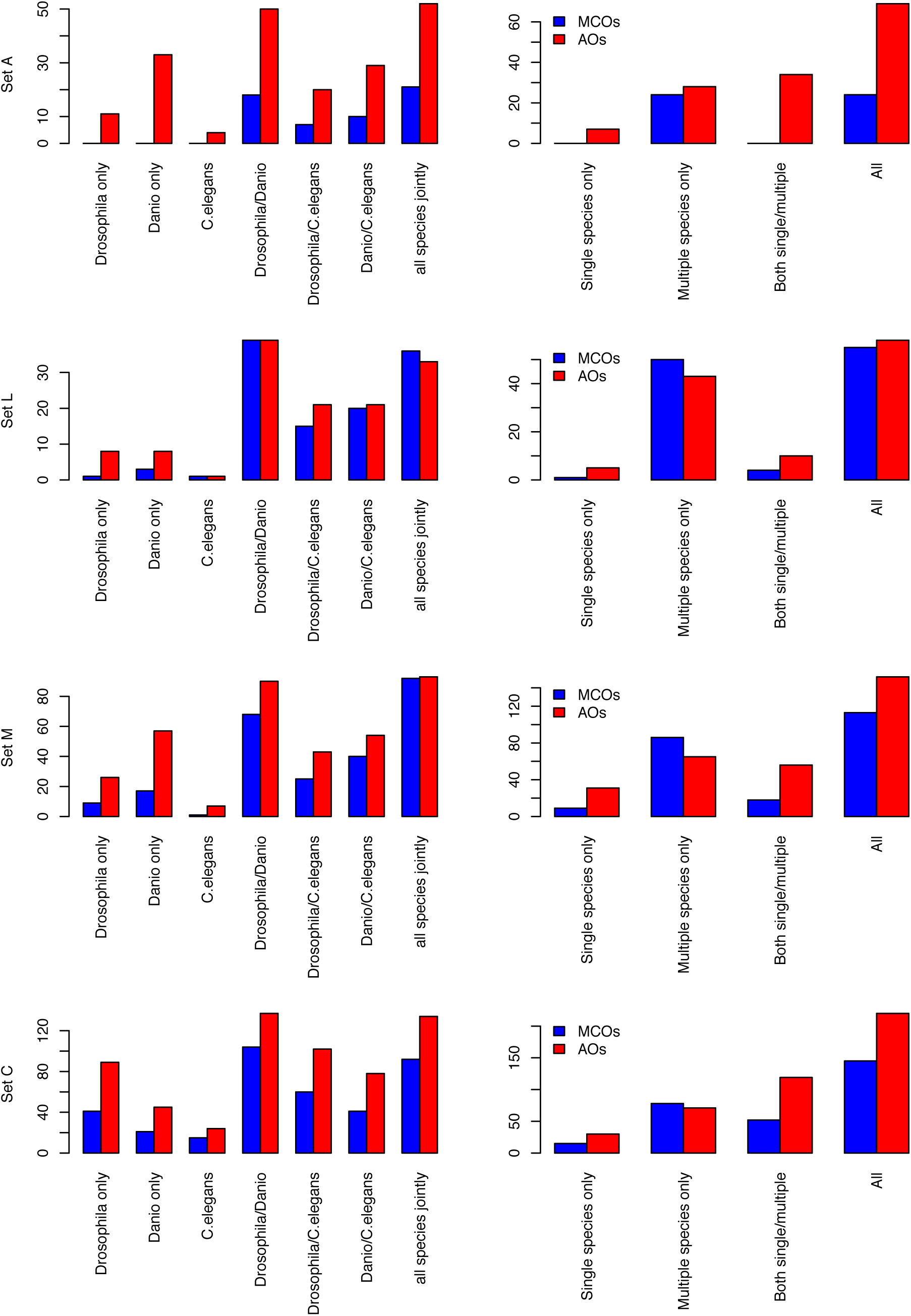
Supplementary Figure related to Figure 3. MSGEA vs. single species enrichment analysis for clusters of orthologous proteins (COPs). Number of significant terms shown for each of the four sets *A*, *L*, *M*, and *C* (cf. Fig 3).

**Figure S2.**
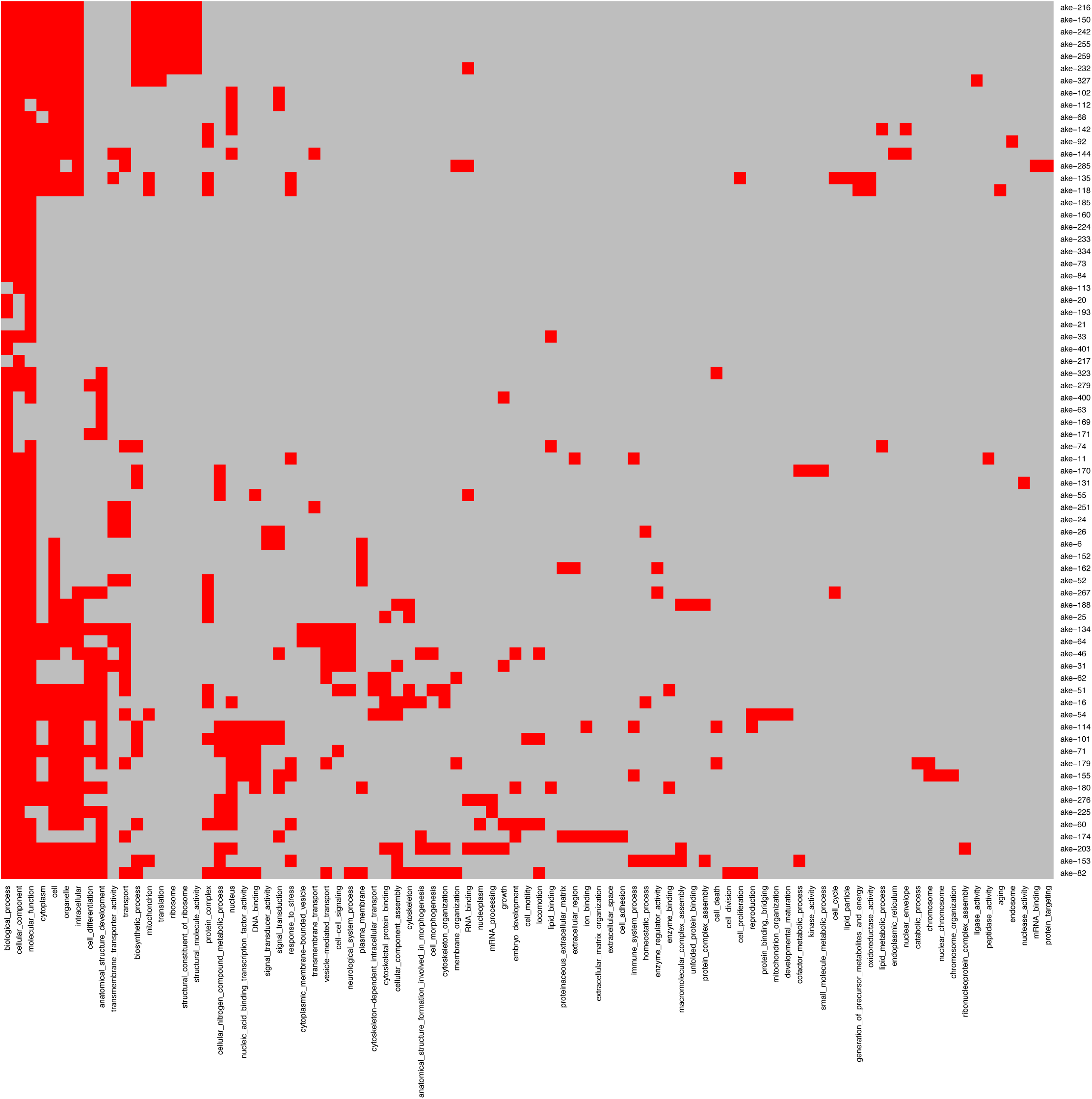
Supplementary Figure related to Figure 4. Absence/presence map of GO-slim terms. *x*-axis: GO-slim terms (including all GO levels); right *y*-axis: internal COP IDs.

**Figure S3.**
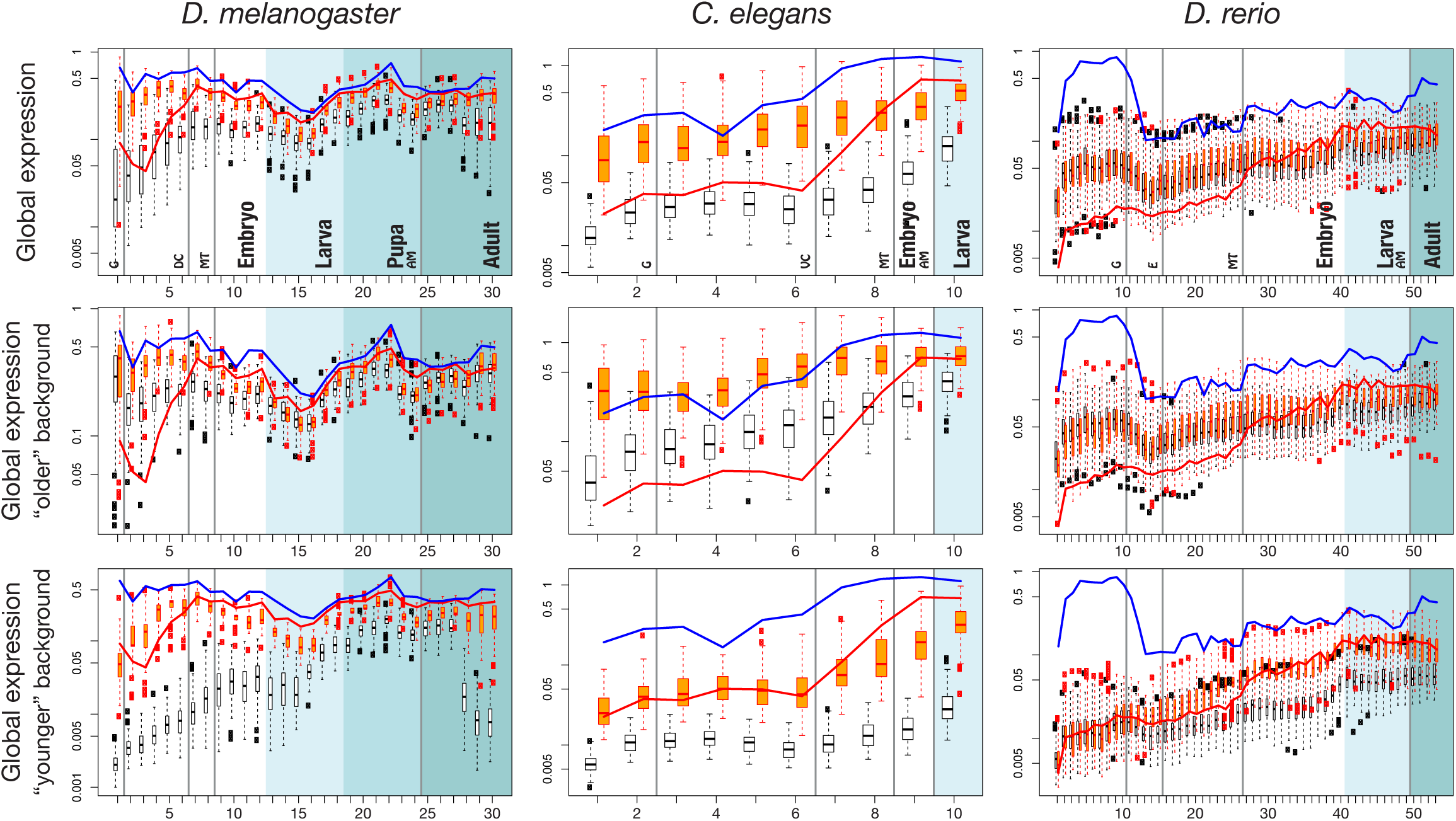
Supplementary Figure related to Figure 5. Expression profiles with background distribution based on medians, instead of means. Color code and other labels as in Fig 2.

**Figure S4.**
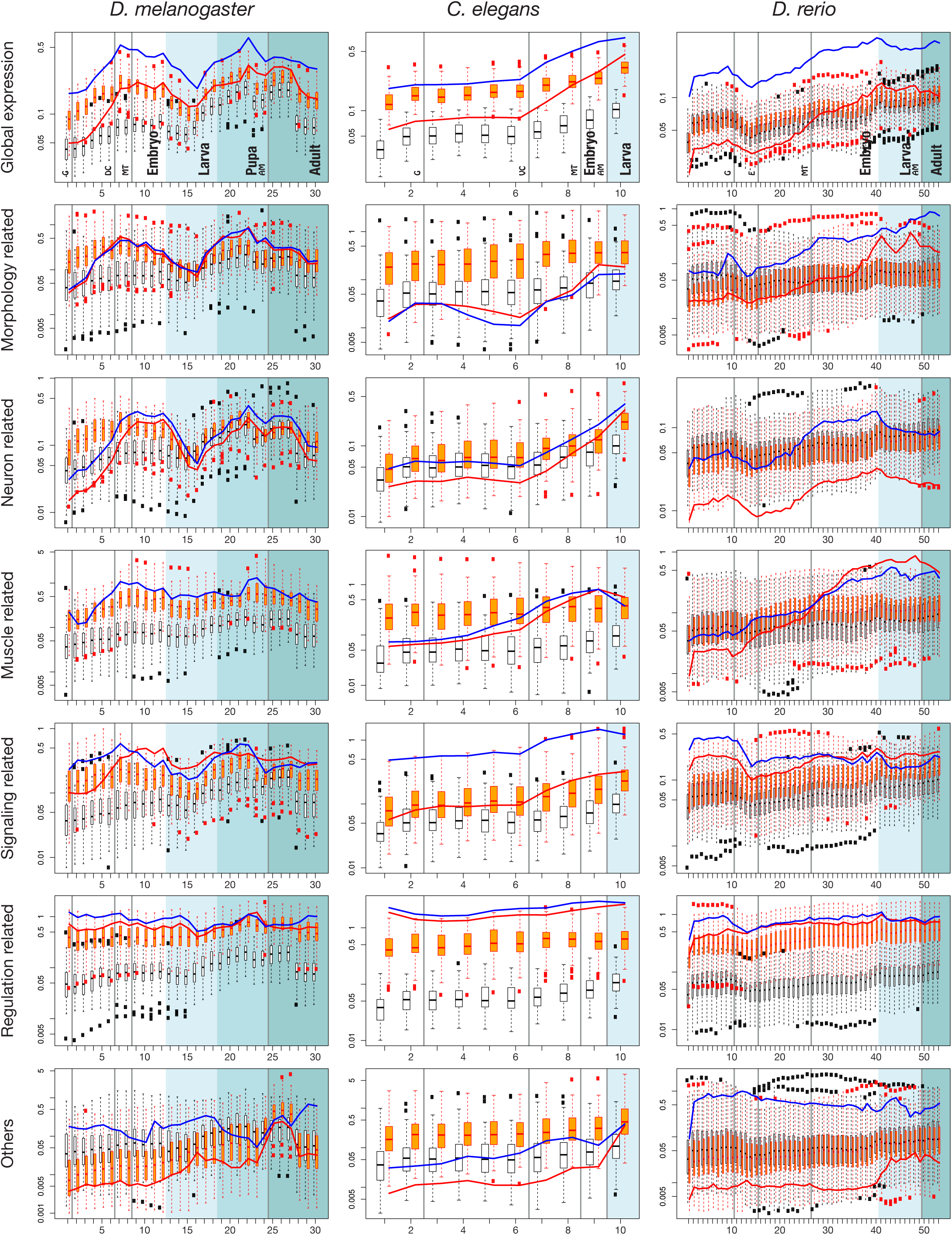
Supplementary Figure related to Figure 5. Expression profiles of set *L*′ genes by functional classes. Background distributions based on mean expression values. Color code and other labels as in Fig 2. *D.m.*, ’muscle’: there are no paralogs; therefore, only the blue line (for MCOs) is displayed.

**Table S1.**
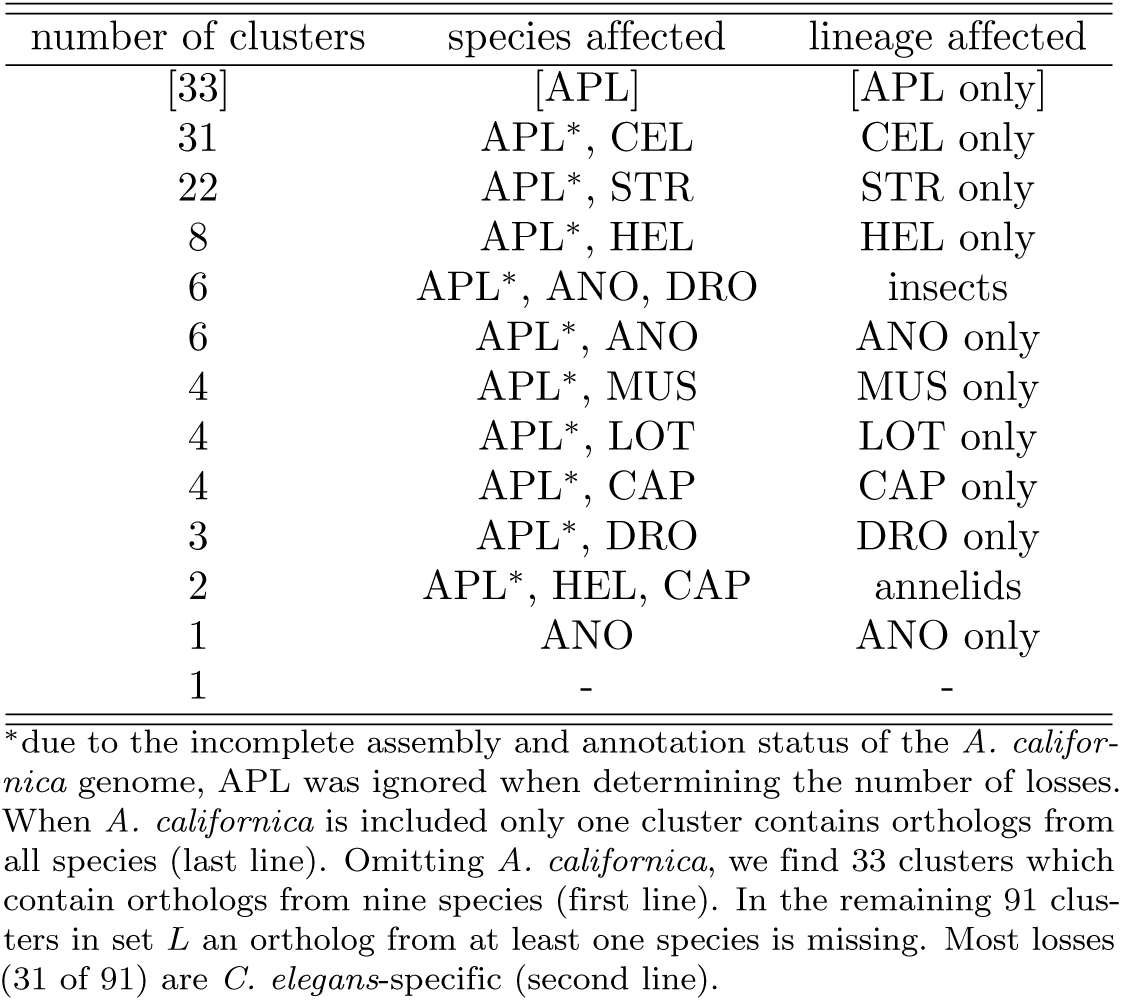
Supplementary Table related to Figure 1. Distribution of losses in set *L*.

| number of clusters | species affected | lineage affected |
| --- | --- | --- |
| [33] | [APL] | [APL only] |
| 31 | APL*, CEL | CEL only |
| 22 | APL*, STR | STR only |
| 8 | APL*, HEL | HEL only |
| 6 | APL*, ANO, DRO | insects |
| 6 | APL*, ANO | ANO only |
| 4 | APL*, MUS | MUS only |
| 4 | APL*, LOT | LOT only |
| 4 | APL*, CAP | CAP only |
| 3 | APL*, DRO | DRO only |
| 2 | APL*, HEL, CAP | annelids |
| 1 | ANO | ANO only |
| 1 | - | - |

\*due to the incomplete assembly and annotation status of the *A. californica* genome, APL was ignored when determining the number of losses. When *A. californica* is included only one cluster contains orthologs from all species (last line). Omitting *A. californica*, we find 33 clusters which contain orthologs from nine species (first line). In the remaining 91 clusters in set *L* an ortholog from at least one species is missing. Most losses (31 of 91) are *C. elegans*-specific (second line).

**Table S2.**
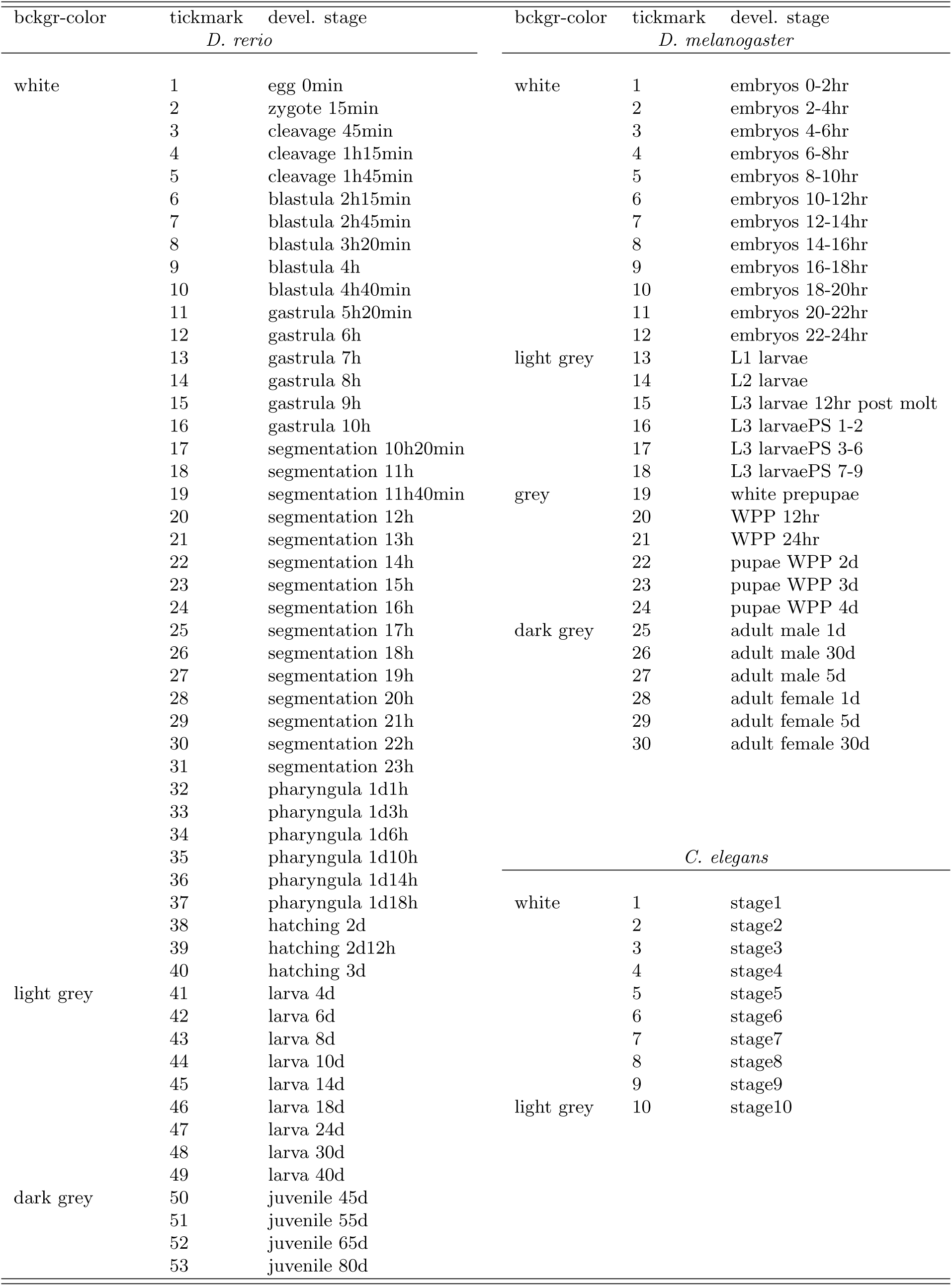
Supplementary Table with annotation of *x*-axis tick marks in Fig 2, Fig S3 and Fig S4 according to [75] (*D. melanogaster*), [124] (*C. elegans*) and [48] (*D. rerio*).

**Table S3.**
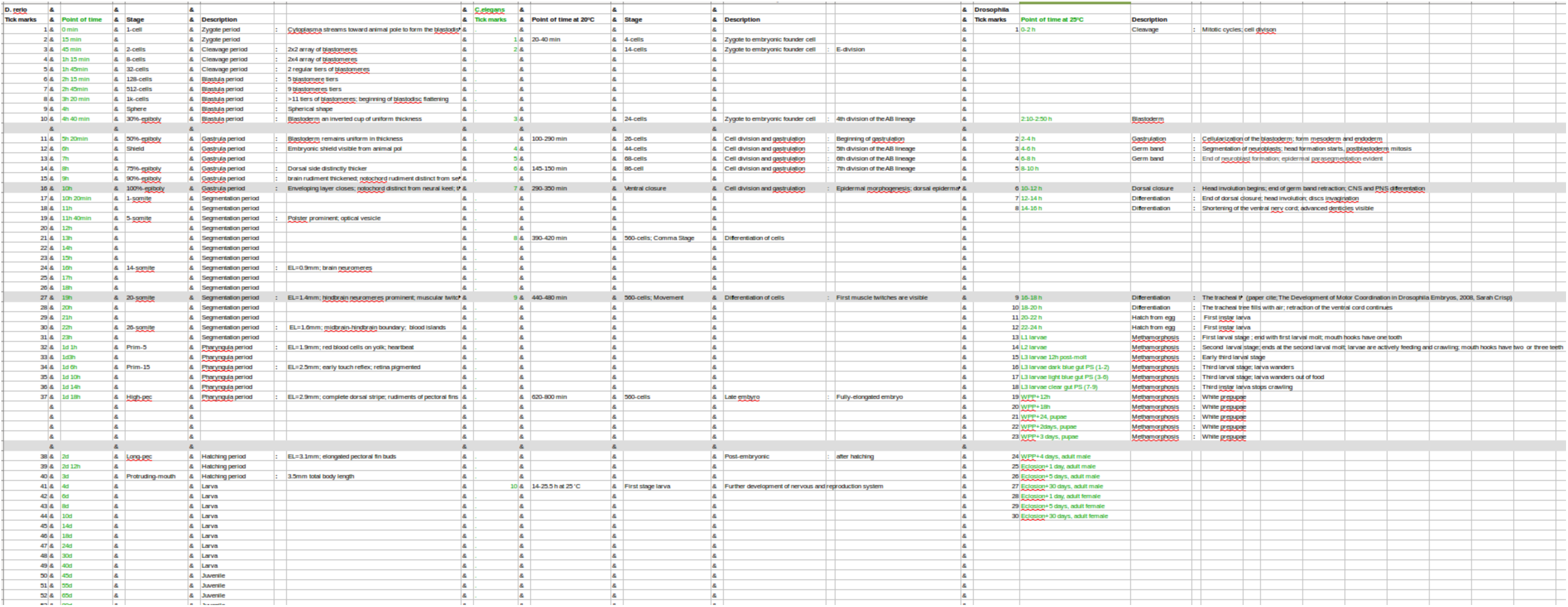
Suppletary Table related to Fig S3 and Fig S4. Comparison of developmental stages among *Drosophila*, *Caenorhabditis*, and *Danio*

**Table S4.**
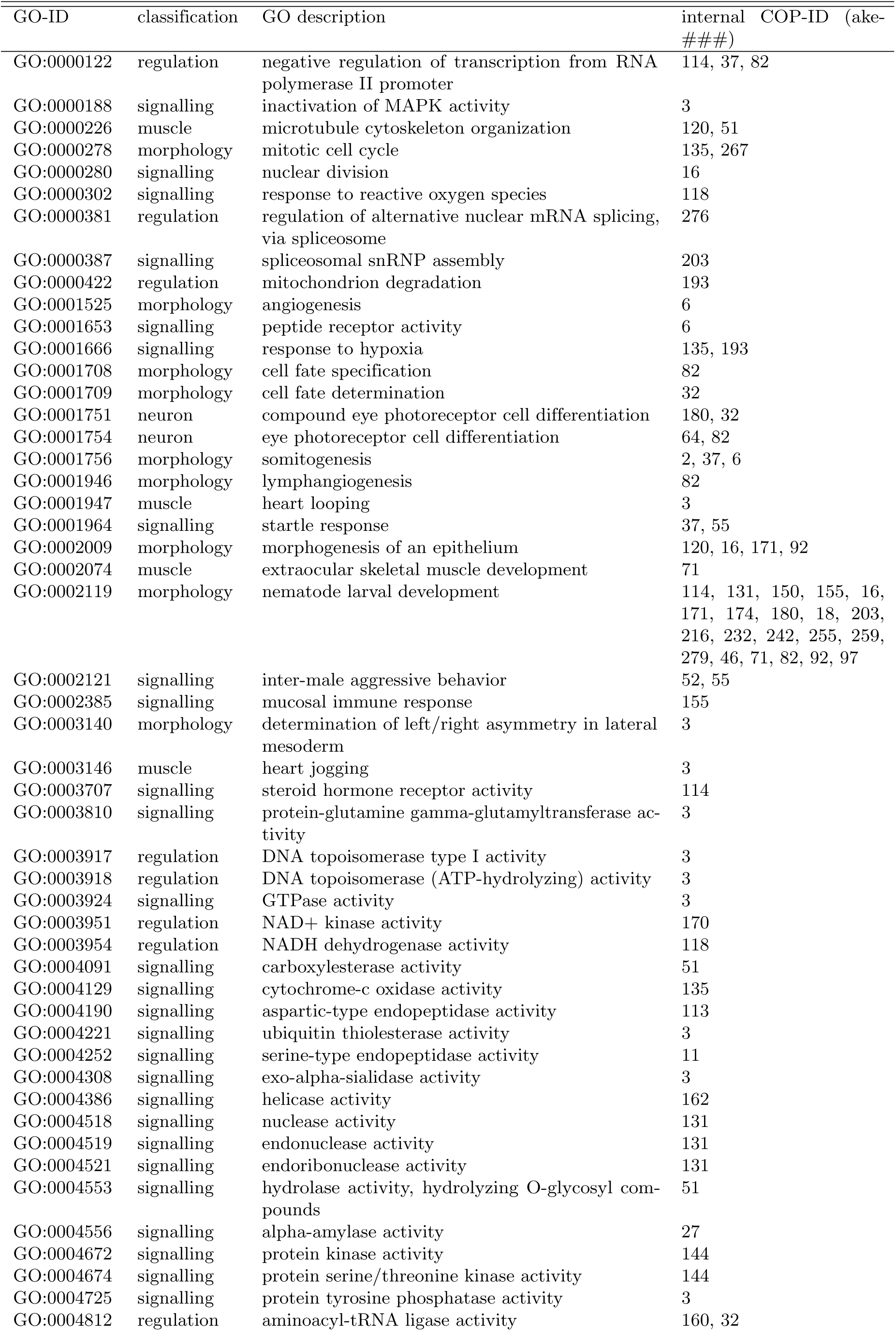
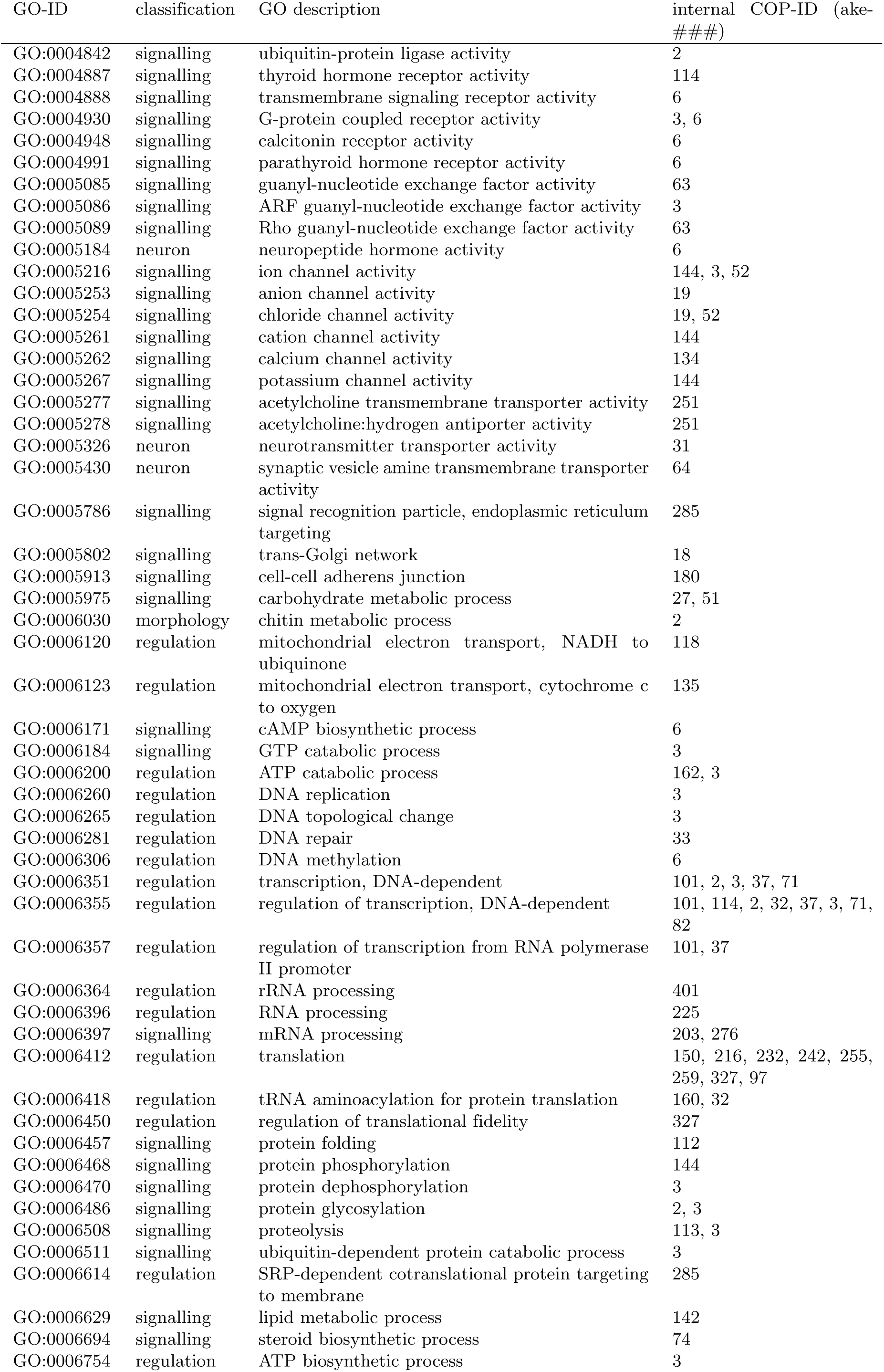
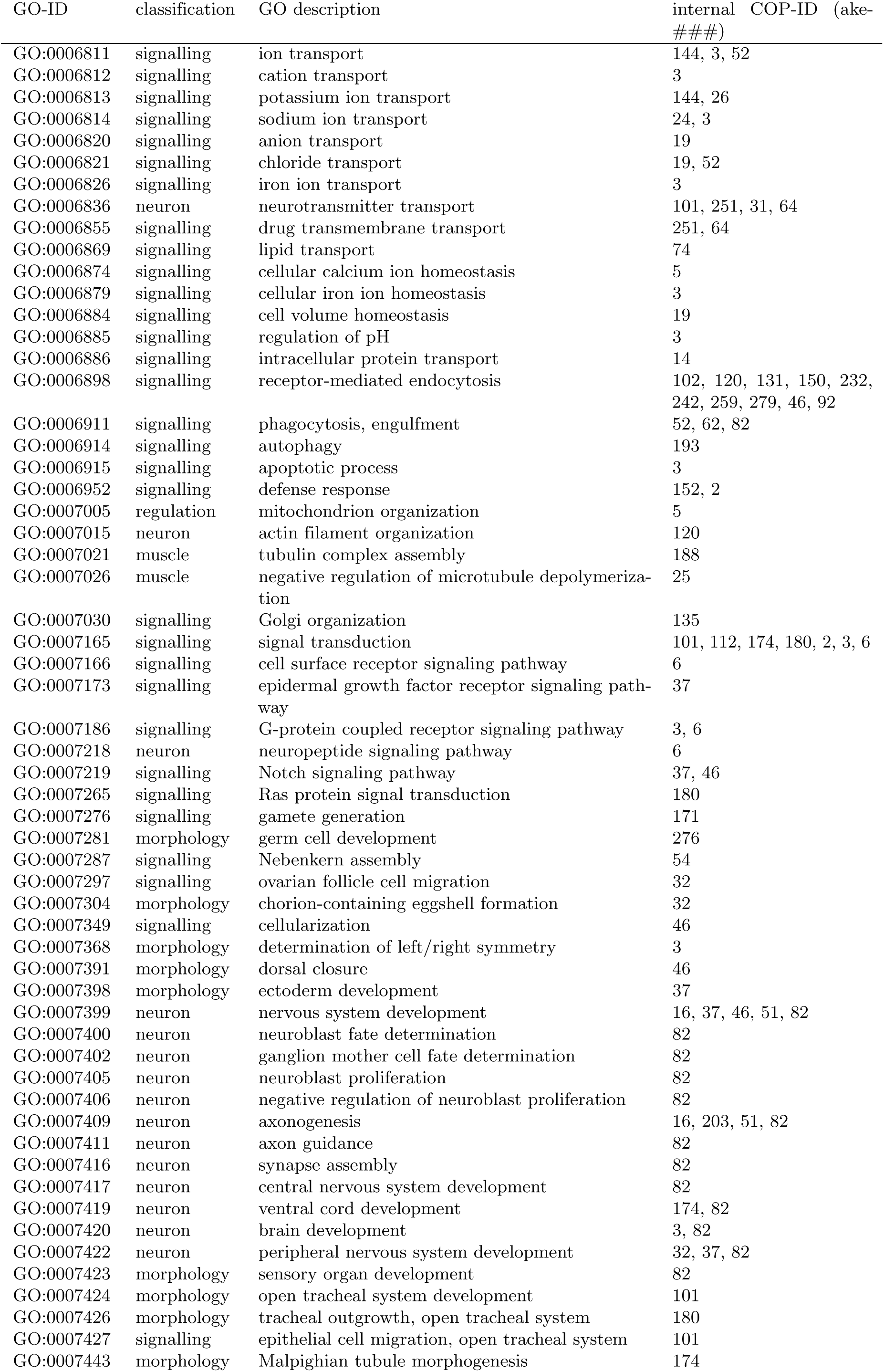
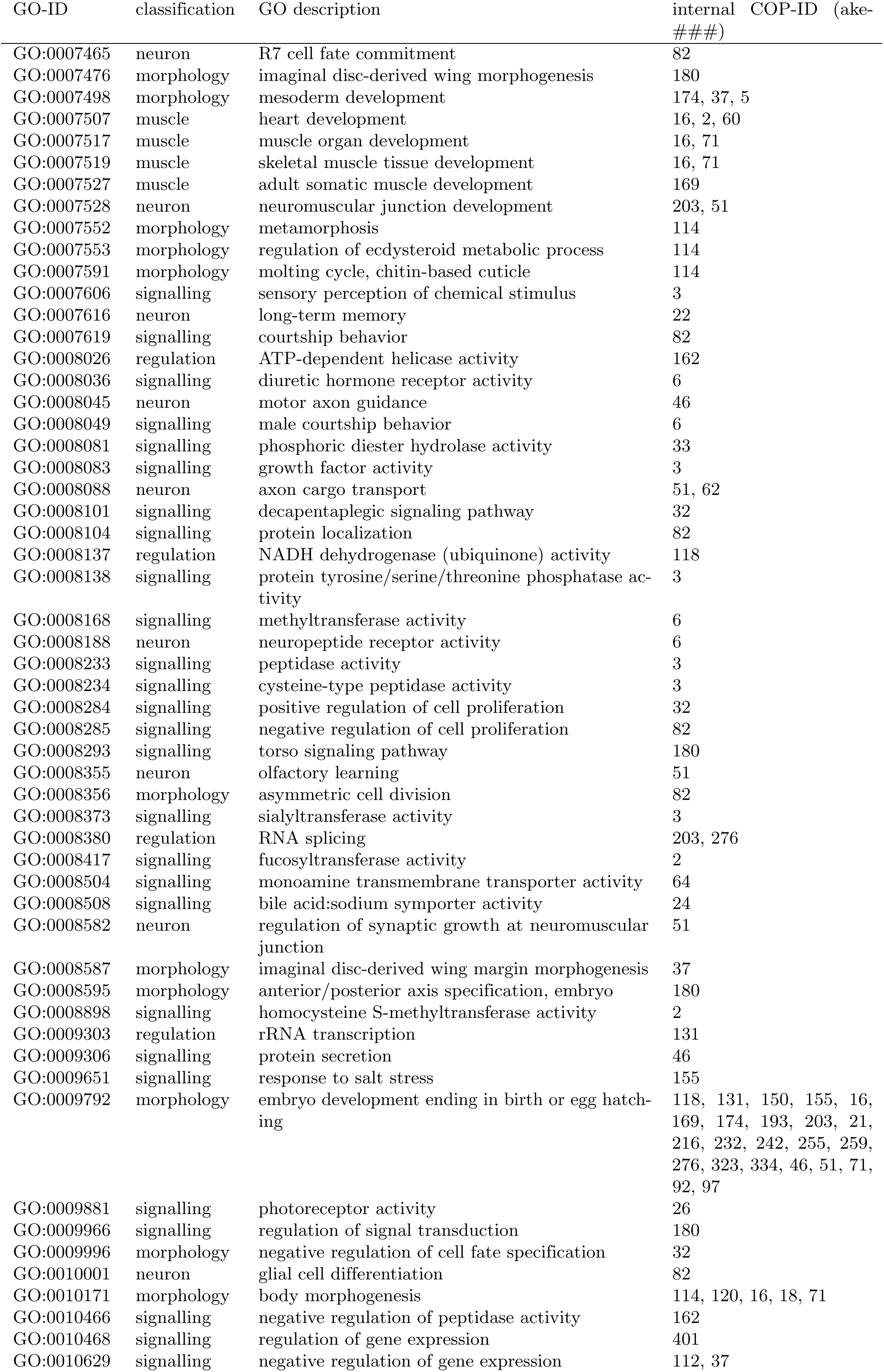
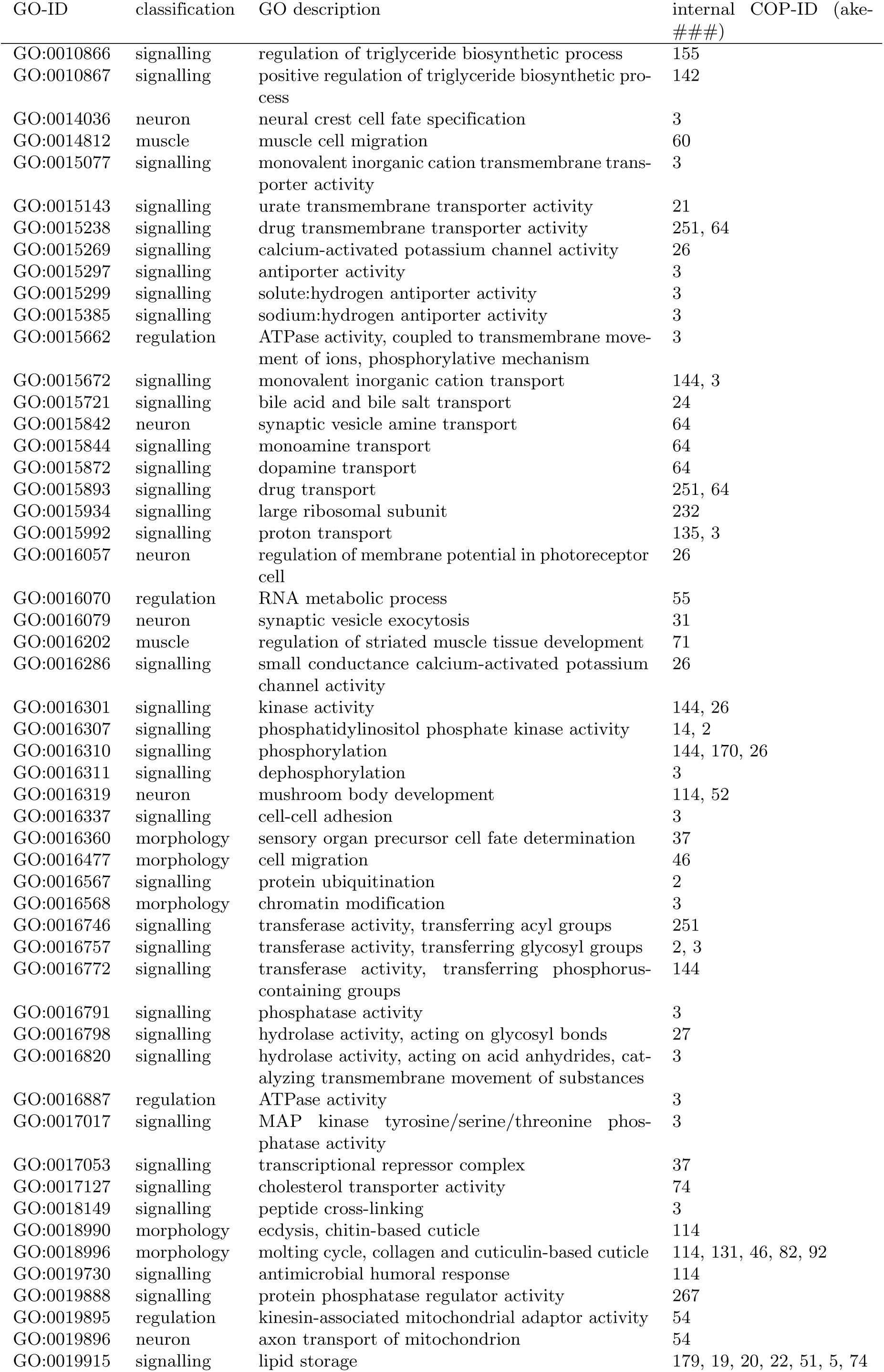
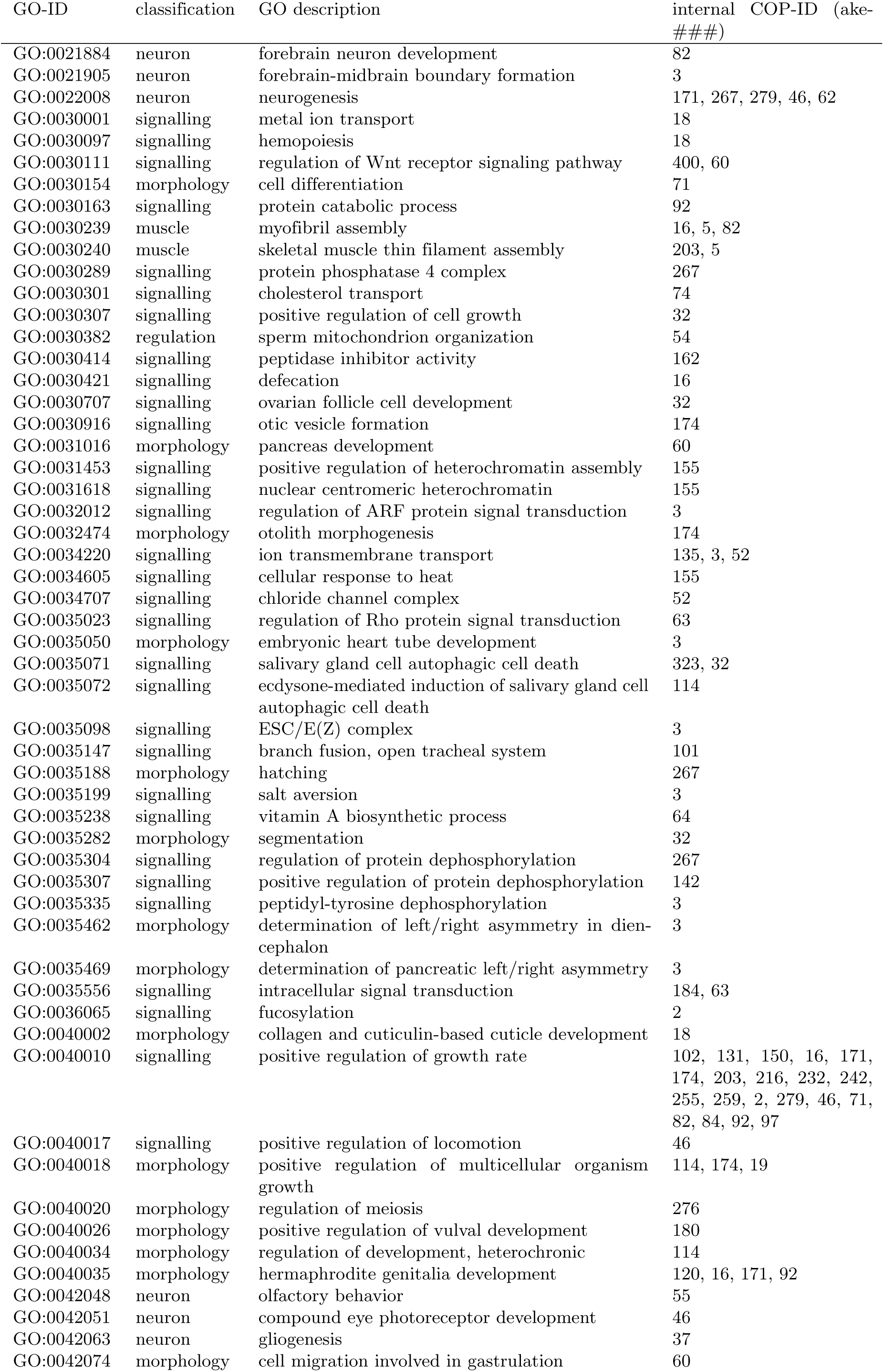
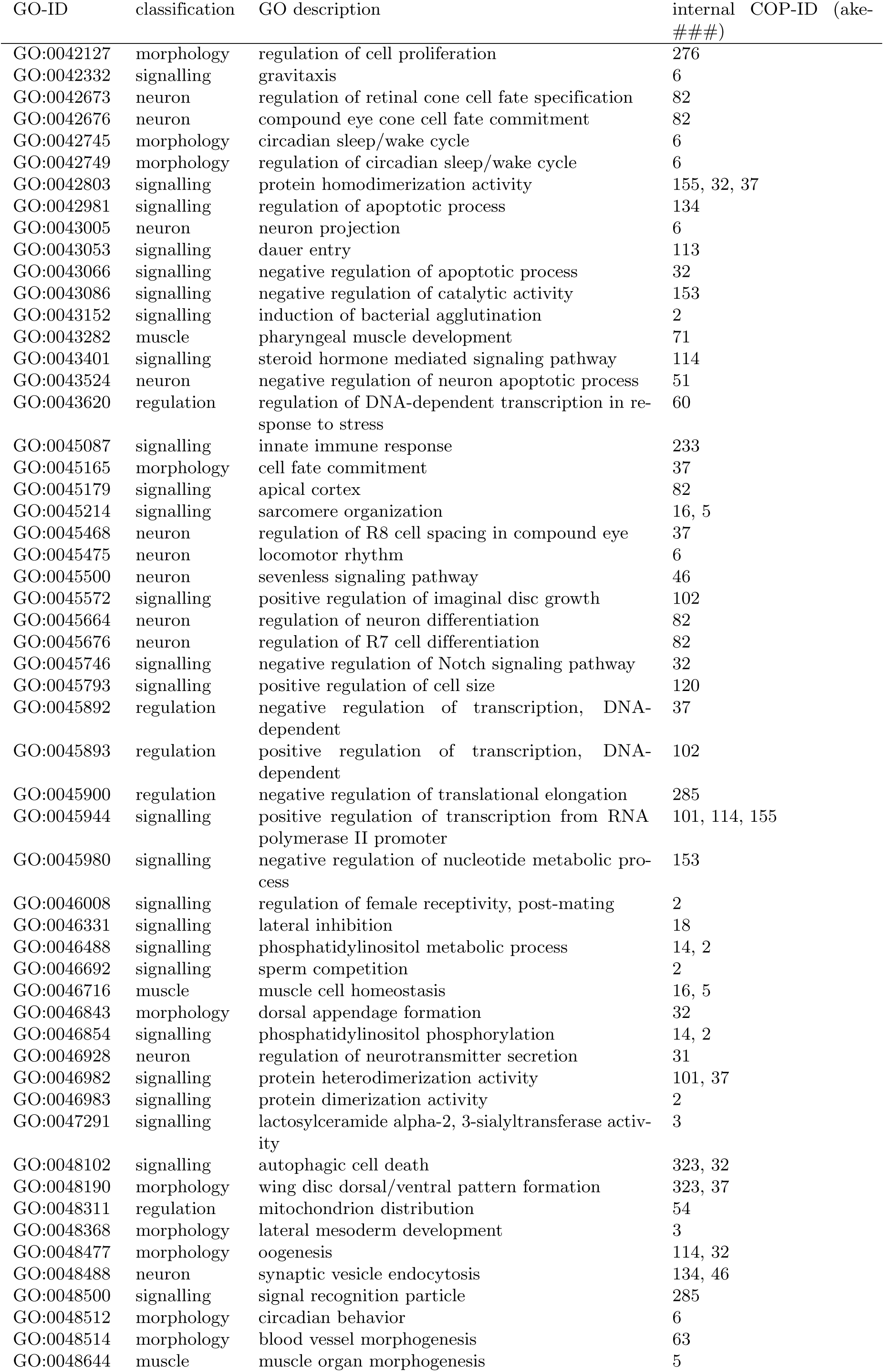
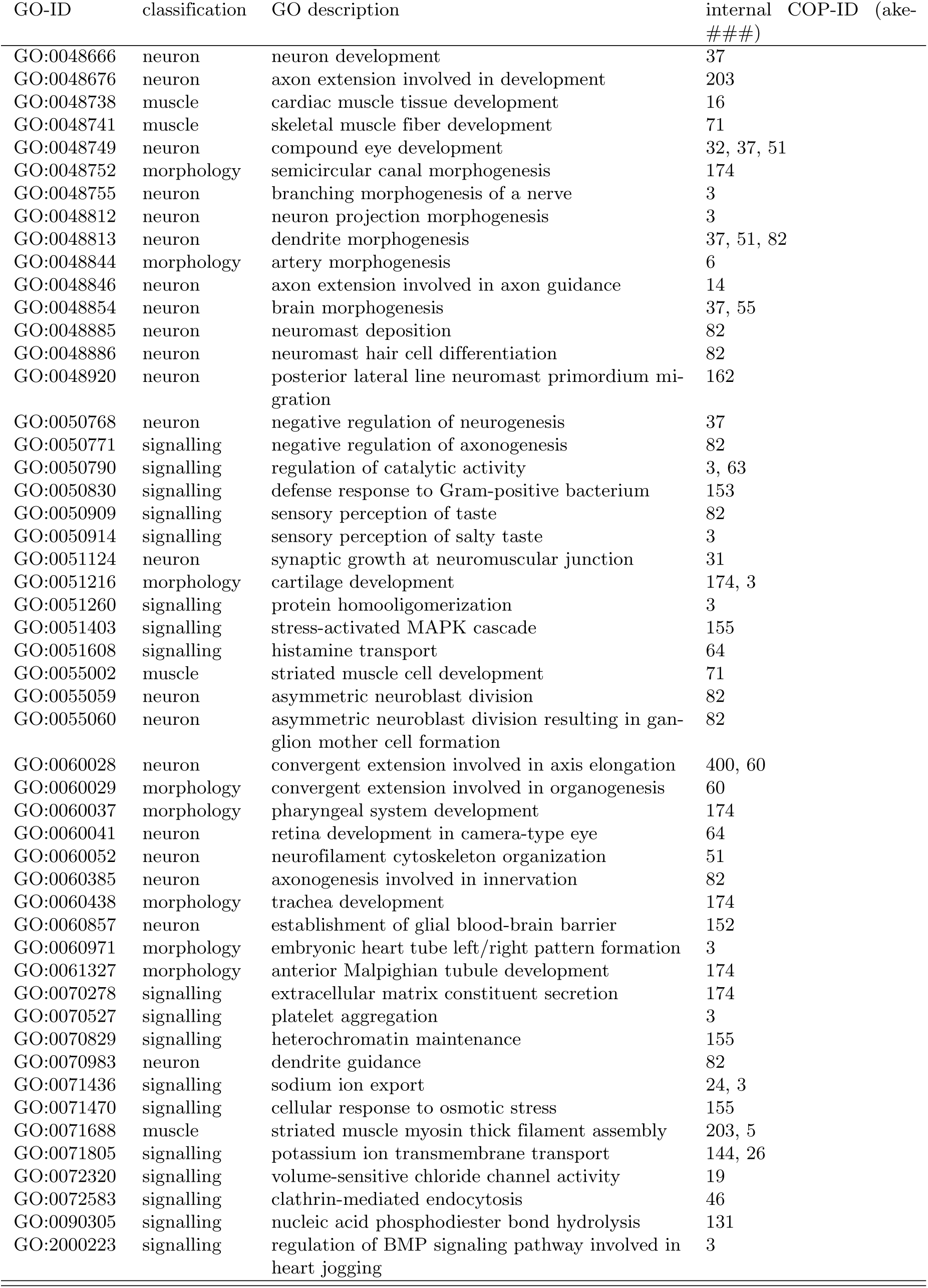
Supplementary Table related to Figures 3 and 4. GO terms (of level ≥4) identified in 85 COPs of set *L*′. COPs ake-68, ake-73, ake-185, ake-217 and ake-224 did not contain GO terms of level ≥4 and do therefore not occur in this table.

| GO-ID | classification | GO description | internal COP-ID (ake-###) |
| --- | --- | --- | --- |
| GO:0000122 | regulation | negative regulation of transcription from RNA polymerase II promoter | 114, 37, 82 |
| GO:0000188 | signalling | inactivation of MAPK activity | 3 |
| GO:0000226 | muscle | microtubule cytoskeleton organization | 120, 51 |
| GO:0000278 | morphology | mitotic cell cycle | 135, 267 |
| GO:0000280 | signalling | nuclear division | 16 |
| GO:0000302 | signalling | response to reactive oxygen species | 118 |
| GO:0000381 | regulation | regulation of alternative nuclear mRNA splicing, via spliceosome | 276 |
| GO:0000387 | signalling | spliceosomal snRNP assembly | 203 |
| GO:0000422 | regulation | mitochondrion degradation | 193 |
| GO:0001525 | morphology | angiogenesis | 6 |
| GO:0001653 | signalling | peptide receptor activity | 6 |
| GO:0001666 | signalling | response to hypoxia | 135, 193 |
| GO:0001708 | morphology | cell fate specification | 82 |
| GO:0001709 | morphology | cell fate determination | 32 |
| GO:0001751 | neuron | compound eye photoreceptor cell differentiation | 180, 32 |
| GO:0001754 | neuron | eye photoreceptor cell differentiation | 64, 82 |
| GO:0001756 | morphology | somitogenesis | 2, 37, 6 |
| GO:0001946 | morphology | lymphangiogenesis | 82 |
| GO:0001947 | muscle | heart looping | 3 |
| GO:0001964 | signalling | startle response | 37, 55 |
| GO:0002009 | morphology | morphogenesis of an epithelium | 120, 16, 171, 92 |
| GO:0002074 | muscle | extraocular skeletal muscle development | 71 |
| GO:0002119 | morphology | nematode larval development | 114, 131, 150, 155, 16, 171, 174, 180, 18, 203, 216, 232, 242, 255, 259, 279, 46, 71, 82, 92, 97 |
| GO:0002121 | signalling | inter-male aggressive behavior | 52, 55 |
| GO:0002385 | signalling | mucosal immune response | 155 |
| GO:0003140 | morphology | determination of left/right asymmetry in lateral mesoderm | 3 |
| GO:0003146 | muscle | heart jogging | 3 |
| GO:0003707 | signalling | steroid hormone receptor activity | 114 |
| GO:0003810 | signalling | protein-glutamine gamma-glutamyltransferase activity | 3 |
| GO:0003917 | regulation | DNA topoisomerase type I activity | 3 |
| GO:0003918 | regulation | DNA topoisomerase (ATP-hydrolyzing) activity | 3 |
| GO:0003924 | signalling | GTPase activity | 3 |
| GO:0003951 | regulation | NAD<sup>+</sup> kinase activity | 170 |
| GO:0003954 | regulation | NADH dehydrogenase activity | 118 |
| GO:0004091 | signalling | carboxylesterase activity | 51 |
| GO:0004129 | signalling | cytochrome-c oxidase activity | 135 |
| GO:0004190 | signalling | aspartic-type endopeptidase activity | 113 |
| GO:0004221 | signalling | ubiquitin thiolesterase activity | 3 |
| GO:0004252 | signalling | serine-type endopeptidase activity | 11 |
| GO:0004308 | signalling | exo-alpha-sialidase activity | 3 |
| GO:0004386 | signalling | helicase activity | 162 |
| GO:0004518 | signalling | nuclease activity | 131 |
| GO:0004519 | signalling | endonuclease activity | 131 |
| GO:0004521 | signalling | endoribonuclease activity | 131 |
| GO:0004553 | signalling | hydrolase activity, hydrolyzing O-glycosyl compounds | 51 |
| GO:0004556 | signalling | alpha-amylase activity | 27 |
| GO:0004672 | signalling | protein kinase activity | 144 |
| GO:0004674 | signalling | protein serine/threonine kinase activity | 144 |
| GO:0004725 | signalling | protein tyrosine phosphatase activity | 3 |
| GO:0004812 | regulation | aminoacyl-tRNA ligase activity | 160, 32 |

| GO-ID | classification | GO description | internal COP-ID (ake-<br>###) |
| --- | --- | --- | --- |
| GO:0004842 | signalling | ubiquitin-protein ligase activity | 2 |
| GO:0004887 | signalling | thyroid hormone receptor activity | 114 |
| GO:0004888 | signalling | transmembrane signaling receptor activity | 6 |
| GO:0004930 | signalling | G-protein coupled receptor activity | 3, 6 |
| GO:0004948 | signalling | calcitonin receptor activity | 6 |
| GO:0004991 | signalling | parathyroid hormone receptor activity | 6 |
| GO:0005085 | signalling | guanyl-nucleotide exchange factor activity | 63 |
| GO:0005086 | signalling | ARF guanyl-nucleotide exchange factor activity | 3 |
| GO:0005089 | signalling | Rho guanyl-nucleotide exchange factor activity | 63 |
| GO:0005184 | neuron | neuropeptide hormone activity | 6 |
| GO:0005216 | signalling | ion channel activity | 144, 3, 52 |
| GO:0005253 | signalling | anion channel activity | 19 |
| GO:0005254 | signalling | chloride channel activity | 19, 52 |
| GO:0005261 | signalling | cation channel activity | 144 |
| GO:0005262 | signalling | calcium channel activity | 134 |
| GO:0005267 | signalling | potassium channel activity | 144 |
| GO:0005277 | signalling | acetylcholine transmembrane transporter activity | 251 |
| GO:0005278 | signalling | acetylcholine:hydrogen antiporter activity | 251 |
| GO:0005326 | neuron | neurotransmitter transporter activity | 31 |
| GO:0005430 | neuron | synaptic vesicle amine transmembrane transporter activity | 64 |
| GO:0005786 | signalling | signal recognition particle, endoplasmic reticulum targeting | 285 |
| GO:0005802 | signalling | trans-Golgi network | 18 |
| GO:0005913 | signalling | cell-cell adherens junction | 180 |
| GO:0005975 | signalling | carbohydrate metabolic process | 27, 51 |
| GO:0006030 | morphology | chitin metabolic process | 2 |
| GO:0006120 | regulation | mitochondrial electron transport, NADH to ubiquinone | 118 |
| GO:0006123 | regulation | mitochondrial electron transport, cytochrome c to oxygen | 135 |
| GO:0006171 | signalling | cAMP biosynthetic process | 6 |
| GO:0006184 | signalling | GTP catabolic process | 3 |
| GO:0006200 | regulation | ATP catabolic process | 162, 3 |
| GO:0006260 | regulation | DNA replication | 3 |
| GO:0006265 | regulation | DNA topological change | 3 |
| GO:0006281 | regulation | DNA repair | 33 |
| GO:0006306 | regulation | DNA methylation | 6 |
| GO:0006351 | regulation | transcription, DNA-dependent | 101, 2, 3, 37, 71 |
| GO:0006355 | regulation | regulation of transcription, DNA-dependent | 101, 114, 2, 32, 37, 3, 71, 82 |
| GO:0006357 | regulation | regulation of transcription from RNA polymerase II promoter | 101, 37 |
| GO:0006364 | regulation | rRNA processing | 401 |
| GO:0006396 | regulation | RNA processing | 225 |
| GO:0006397 | signalling | mRNA processing | 203, 276 |
| GO:0006412 | regulation | translation | 150, 216, 232, 242, 255, 259, 327, 97 |
| GO:0006418 | regulation | tRNA aminoacylation for protein translation | 160, 32 |
| GO:0006450 | regulation | regulation of translational fidelity | 327 |
| GO:0006457 | signalling | protein folding | 112 |
| GO:0006468 | signalling | protein phosphorylation | 144 |
| GO:0006470 | signalling | protein dephosphorylation | 3 |
| GO:0006486 | signalling | protein glycosylation | 2, 3 |
| GO:0006508 | signalling | proteolysis | 113, 3 |
| GO:0006511 | signalling | ubiquitin-dependent protein catabolic process | 3 |
| GO:0006614 | regulation | SRP-dependent cotranslational protein targeting to membrane | 285 |
| GO:0006629 | signalling | lipid metabolic process | 142 |
| GO:0006694 | signalling | steroid biosynthetic process | 74 |
| GO:0006754 | regulation | ATP biosynthetic process | 3 |
| GO:0006811 | signalling | ion transport | 144, 3, 52 |
| GO:0006812 | signalling | cation transport | 3 |
| GO:0006813 | signalling | potassium ion transport | 144, 26 |
| GO:0006814 | signalling | sodium ion transport | 24, 3 |
| GO:0006820 | signalling | anion transport | 19 |
| GO:0006821 | signalling | chloride transport | 19, 52 |
| GO:0006826 | signalling | iron ion transport | 3 |
| GO:0006836 | neuron | neurotransmitter transport | 101, 251, 31, 64 |
| GO:0006855 | signalling | drug transmembrane transport | 251, 64 |
| GO:0006869 | signalling | lipid transport | 74 |
| GO:0006874 | signalling | cellular calcium ion homeostasis | 5 |
| GO:0006879 | signalling | cellular iron ion homeostasis | 3 |
| GO:0006884 | signalling | cell volume homeostasis | 19 |
| GO:0006885 | signalling | regulation of pH | 3 |
| GO:0006886 | signalling | intracellular protein transport | 14 |
| GO:0006898 | signalling | receptor-mediated endocytosis | 102, 120, 131, 150, 232,<br>242, 259, 279, 46, 92 |
| GO:0006911 | signalling | phagocytosis, engulfment | 52, 62, 82 |
| GO:0006914 | signalling | autophagy | 193 |
| GO:0006915 | signalling | apoptotic process | 3 |
| GO:0006952 | signalling | defense response | 152, 2 |
| GO:0007005 | regulation | mitochondrion organization | 5 |
| GO:0007015 | neuron | actin filament organization | 120 |
| GO:0007021 | muscle | tubulin complex assembly | 188 |
| GO:0007026 | muscle | negative regulation of microtubule depolymeriza-<br>tion | 25 |
| GO:0007030 | signalling | Golgi organization | 135 |
| GO:0007165 | signalling | signal transduction | 101, 112, 174, 180, 2, 3, 6 |
| GO:0007166 | signalling | cell surface receptor signaling pathway | 6 |
| GO:0007173 | signalling | epidermal growth factor receptor signaling path-<br>way | 37 |
| GO:0007186 | signalling | G-protein coupled receptor signaling pathway | 3, 6 |
| GO:0007218 | neuron | neuropeptide signaling pathway | 6 |
| GO:0007219 | signalling | Notch signaling pathway | 37, 46 |
| GO:0007265 | signalling | Ras protein signal transduction | 180 |
| GO:0007276 | signalling | gamete generation | 171 |
| GO:0007281 | morphology | germ cell development | 276 |
| GO:0007287 | signalling | Nebenkern assembly | 54 |
| GO:0007297 | signalling | ovarian follicle cell migration | 32 |
| GO:0007304 | morphology | chorion-containing eggshell formation | 32 |
| GO:0007349 | signalling | cellularization | 46 |
| GO:0007368 | morphology | determination of left/right symmetry | 3 |
| GO:0007391 | morphology | dorsal closure | 46 |
| GO:0007398 | morphology | ectoderm development | 37 |
| GO:0007399 | neuron | nervous system development | 16, 37, 46, 51, 82 |
| GO:0007400 | neuron | neuroblast fate determination | 82 |
| GO:0007402 | neuron | ganglion mother cell fate determination | 82 |
| GO:0007405 | neuron | neuroblast proliferation | 82 |
| GO:0007406 | neuron | negative regulation of neuroblast proliferation | 82 |
| GO:0007409 | neuron | axonogenesis | 16, 203, 51, 82 |
| GO:0007411 | neuron | axon guidance | 82 |
| GO:0007416 | neuron | synapse assembly | 82 |
| GO:0007417 | neuron | central nervous system development | 82 |
| GO:0007419 | neuron | ventral cord development | 174, 82 |
| GO:0007420 | neuron | brain development | 3, 82 |
| GO:0007422 | neuron | peripheral nervous system development | 32, 37, 82 |
| GO:0007423 | morphology | sensory organ development | 82 |
| GO:0007424 | morphology | open tracheal system development | 101 |
| GO:0007426 | morphology | tracheal outgrowth, open tracheal system | 180 |
| GO:0007427 | signalling | epithelial cell migration, open tracheal system | 101 |
| GO:0007443 | morphology | Malpighian tubule morphogenesis | 174 |
| GO:0007465 | neuron | R7 cell fate commitment | 82 |
| GO:0007476 | morphology | imaginal disc-derived wing morphogenesis | 180 |
| GO:0007498 | morphology | mesoderm development | 174, 37, 5 |
| GO:0007507 | muscle | heart development | 16, 2, 60 |
| GO:0007517 | muscle | muscle organ development | 16, 71 |
| GO:0007519 | muscle | skeletal muscle tissue development | 16, 71 |
| GO:0007527 | muscle | adult somatic muscle development | 169 |
| GO:0007528 | neuron | neuromuscular junction development | 203, 51 |
| GO:0007552 | morphology | metamorphosis | 114 |
| GO:0007553 | morphology | regulation of ecdysteroid metabolic process | 114 |
| GO:0007591 | morphology | molting cycle, chitin-based cuticle | 114 |
| GO:0007606 | signalling | sensory perception of chemical stimulus | 3 |
| GO:0007616 | neuron | long-term memory | 22 |
| GO:0007619 | signalling | courtship behavior | 82 |
| GO:0008026 | regulation | ATP-dependent helicase activity | 162 |
| GO:0008036 | signalling | diuretic hormone receptor activity | 6 |
| GO:0008045 | neuron | motor axon guidance | 46 |
| GO:0008049 | signalling | male courtship behavior | 6 |
| GO:0008081 | signalling | phosphoric diester hydrolase activity | 33 |
| GO:0008083 | signalling | growth factor activity | 3 |
| GO:0008088 | neuron | axon cargo transport | 51, 62 |
| GO:0008101 | signalling | decapentaplegic signaling pathway | 32 |
| GO:0008104 | signalling | protein localization | 82 |
| GO:0008137 | regulation | NADH dehydrogenase (ubiquinone) activity | 118 |
| GO:0008138 | signalling | protein tyrosine/serine/threonine phosphatase activity | 3 |
| GO:0008168 | signalling | methyltransferase activity | 6 |
| GO:0008188 | neuron | neuropeptide receptor activity | 6 |
| GO:0008233 | signalling | peptidase activity | 3 |
| GO:0008234 | signalling | cysteine-type peptidase activity | 3 |
| GO:0008284 | signalling | positive regulation of cell proliferation | 32 |
| GO:0008285 | signalling | negative regulation of cell proliferation | 82 |
| GO:0008293 | signalling | torso signaling pathway | 180 |
| GO:0008355 | neuron | olfactory learning | 51 |
| GO:0008356 | morphology | asymmetric cell division | 82 |
| GO:0008373 | signalling | sialyltransferase activity | 3 |
| GO:0008380 | regulation | RNA splicing | 203, 276 |
| GO:0008417 | signalling | fucosyltransferase activity | 2 |
| GO:0008504 | signalling | monoamine transmembrane transporter activity | 64 |
| GO:0008508 | signalling | bile acid:sodium symporter activity | 24 |
| GO:0008582 | neuron | regulation of synaptic growth at neuromuscular junction | 51 |
| GO:0008587 | morphology | imaginal disc-derived wing margin morphogenesis | 37 |
| GO:0008595 | morphology | anterior/posterior axis specification, embryo | 180 |
| GO:0008898 | signalling | homocysteine S-methyltransferase activity | 2 |
| GO:0009303 | regulation | rRNA transcription | 131 |
| GO:0009306 | signalling | protein secretion | 46 |
| GO:0009651 | signalling | response to salt stress | 155 |
| GO:0009792 | morphology | embryo development ending in birth or egg hatching | 118, 131, 150, 155, 16, 169, 174, 193, 203, 21, 216, 232, 242, 255, 259, 276, 323, 334, 46, 51, 71, 92, 97 |
| GO:0009881 | signalling | photoreceptor activity | 26 |
| GO:0009966 | signalling | regulation of signal transduction | 180 |
| GO:0009996 | morphology | negative regulation of cell fate specification | 32 |
| GO:0010001 | neuron | glial cell differentiation | 82 |
| GO:0010171 | morphology | body morphogenesis | 114, 120, 16, 18, 71 |
| GO:0010466 | signalling | negative regulation of peptidase activity | 162 |
| GO:0010468 | signalling | regulation of gene expression | 401 |
| GO:0010629 | signalling | negative regulation of gene expression | 112, 37 |
| GO:0010866 | signalling | regulation of triglyceride biosynthetic process | 155 |
| GO:0010867 | signalling | positive regulation of triglyceride biosynthetic process | 142 |
| GO:0014036 | neuron | neural crest cell fate specification | 3 |
| GO:0014812 | muscle | muscle cell migration | 60 |
| GO:0015077 | signalling | monovalent inorganic cation transmembrane transporter activity | 3 |
| GO:0015143 | signalling | urate transmembrane transporter activity | 21 |
| GO:0015238 | signalling | drug transmembrane transporter activity | 251, 64 |
| GO:0015269 | signalling | calcium-activated potassium channel activity | 26 |
| GO:0015297 | signalling | antiporter activity | 3 |
| GO:0015299 | signalling | solute:hydrogen antiporter activity | 3 |
| GO:0015385 | signalling | sodium:hydrogen antiporter activity | 3 |
| GO:0015662 | regulation | ATPase activity, coupled to transmembrane movement of ions, phosphorylative mechanism | 3 |
| GO:0015672 | signalling | monovalent inorganic cation transport | 144, 3 |
| GO:0015721 | signalling | bile acid and bile salt transport | 24 |
| GO:0015842 | neuron | synaptic vesicle amine transport | 64 |
| GO:0015844 | signalling | monoamine transport | 64 |
| GO:0015872 | signalling | dopamine transport | 64 |
| GO:0015893 | signalling | drug transport | 251, 64 |
| GO:0015934 | signalling | large ribosomal subunit | 232 |
| GO:0015992 | signalling | proton transport | 135, 3 |
| GO:0016057 | neuron | regulation of membrane potential in photoreceptor cell | 26 |
| GO:0016070 | regulation | RNA metabolic process | 55 |
| GO:0016079 | neuron | synaptic vesicle exocytosis | 31 |
| GO:0016202 | muscle | regulation of striated muscle tissue development | 71 |
| GO:0016286 | signalling | small conductance calcium-activated potassium channel activity | 26 |
| GO:0016301 | signalling | kinase activity | 144, 26 |
| GO:0016307 | signalling | phosphatidylinositol phosphate kinase activity | 14, 2 |
| GO:0016310 | signalling | phosphorylation | 144, 170, 26 |
| GO:0016311 | signalling | dephosphorylation | 3 |
| GO:0016319 | neuron | mushroom body development | 114, 52 |
| GO:0016337 | signalling | cell-cell adhesion | 3 |
| GO:0016360 | morphology | sensory organ precursor cell fate determination | 37 |
| GO:0016477 | morphology | cell migration | 46 |
| GO:0016567 | signalling | protein ubiquitination | 2 |
| GO:0016568 | morphology | chromatin modification | 3 |
| GO:0016746 | signalling | transferase activity, transferring acyl groups | 251 |
| GO:0016757 | signalling | transferase activity, transferring glycosyl groups | 2, 3 |
| GO:0016772 | signalling | transferase activity, transferring phosphorus-containing groups | 144 |
| GO:0016791 | signalling | phosphatase activity | 3 |
| GO:0016798 | signalling | hydrolase activity, acting on glycosyl bonds | 27 |
| GO:0016820 | signalling | hydrolase activity, acting on acid anhydrides, catalyzing transmembrane movement of substances | 3 |
| GO:0016887 | regulation | ATPase activity | 3 |
| GO:0017017 | signalling | MAP kinase tyrosine/serine/threonine phosphatase activity | 3 |
| GO:0017053 | signalling | transcriptional repressor complex | 37 |
| GO:0017127 | signalling | cholesterol transporter activity | 74 |
| GO:0018149 | signalling | peptide cross-linking | 3 |
| GO:0018990 | morphology | ecdysis, chitin-based cuticle | 114 |
| GO:0018996 | morphology | molting cycle, collagen and cuticulin-based cuticle | 114, 131, 46, 82, 92 |
| GO:0019730 | signalling | antimicrobial humoral response | 114 |
| GO:0019888 | signalling | protein phosphatase regulator activity | 267 |
| GO:0019895 | regulation | kinesin-associated mitochondrial adaptor activity | 54 |
| GO:0019896 | neuron | axon transport of mitochondrion | 54 |
| GO:0019915 | signalling | lipid storage | 179, 19, 20, 22, 51, 5, 74 |

... Table S4 continued
| GO-ID | classification | GO description | internal COP-ID (ake-<br>###) |
| --- | --- | --- | --- |
| GO:0021884 | neuron | forebrain neuron development | 82 |
| GO:0021905 | neuron | forebrain-midbrain boundary formation | 3 |
| GO:0022008 | neuron | neurogenesis | 171, 267, 279, 46, 62 |
| GO:0030001 | signalling | metal ion transport | 18 |
| GO:0030097 | signalling | hemopoiesis | 18 |
| GO:0030111 | signalling | regulation of Wnt receptor signaling pathway | 400, 60 |
| GO:0030154 | morphology | cell differentiation | 71 |
| GO:0030163 | signalling | protein catabolic process | 92 |
| GO:0030239 | muscle | myofibril assembly | 16, 5, 82 |
| GO:0030240 | muscle | skeletal muscle thin filament assembly | 203, 5 |
| GO:0030289 | signalling | protein phosphatase 4 complex | 267 |
| GO:0030301 | signalling | cholesterol transport | 74 |
| GO:0030307 | signalling | positive regulation of cell growth | 32 |
| GO:0030382 | regulation | sperm mitochondrion organization | 54 |
| GO:0030414 | signalling | peptidase inhibitor activity | 162 |
| GO:0030421 | signalling | defecation | 16 |
| GO:0030707 | signalling | ovarian follicle cell development | 32 |
| GO:0030916 | signalling | otic vesicle formation | 174 |
| GO:0031016 | morphology | pancreas development | 60 |
| GO:0031453 | signalling | positive regulation of heterochromatin assembly | 155 |
| GO:0031618 | signalling | nuclear centromeric heterochromatin | 155 |
| GO:0032012 | signalling | regulation of ARF protein signal transduction | 3 |
| GO:0032474 | morphology | otolith morphogenesis | 174 |
| GO:0034220 | signalling | ion transmembrane transport | 135, 3, 52 |
| GO:0034605 | signalling | cellular response to heat | 155 |
| GO:0034707 | signalling | chloride channel complex | 52 |
| GO:0035023 | signalling | regulation of Rho protein signal transduction | 63 |
| GO:0035050 | morphology | embryonic heart tube development | 3 |
| GO:0035071 | signalling | salivary gland cell autophagic cell death | 323, 32 |
| GO:0035072 | signalling | ecdysone-mediated induction of salivary gland cell<br>autophagic cell death | 114 |
| GO:0035098 | signalling | ESC/E(Z) complex | 3 |
| GO:0035147 | signalling | branch fusion, open tracheal system | 101 |
| GO:0035188 | morphology | hatching | 267 |
| GO:0035199 | signalling | salt aversion | 3 |
| GO:0035238 | signalling | vitamin A biosynthetic process | 64 |
| GO:0035282 | morphology | segmentation | 32 |
| GO:0035304 | signalling | regulation of protein dephosphorylation | 267 |
| GO:0035307 | signalling | positive regulation of protein dephosphorylation | 142 |
| GO:0035335 | signalling | peptidyl-tyrosine dephosphorylation | 3 |
| GO:0035462 | morphology | determination of left/right asymmetry in dien-<br>cephalon | 3 |
| GO:0035469 | morphology | determination of pancreatic left/right asymmetry | 3 |
| GO:0035556 | signalling | intracellular signal transduction | 184, 63 |
| GO:0036065 | signalling | fucosylation | 2 |
| GO:0040002 | morphology | collagen and cuticulin-based cuticle development | 18 |
| GO:0040010 | signalling | positive regulation of growth rate | 102, 131, 150, 16, 171,<br>174, 203, 216, 232, 242,<br>255, 259, 2, 279, 46, 71,<br>82, 84, 92, 97 |
| GO:0040017 | signalling | positive regulation of locomotion | 46 |
| GO:0040018 | morphology | positive regulation of multicellular organism<br>growth | 114, 174, 19 |
| GO:0040020 | morphology | regulation of meiosis | 276 |
| GO:0040026 | morphology | positive regulation of vulval development | 180 |
| GO:0040034 | morphology | regulation of development, heterochronic | 114 |
| GO:0040035 | morphology | hermaphrodite genitalia development | 120, 16, 171, 92 |
| GO:0042048 | neuron | olfactory behavior | 55 |
| GO:0042051 | neuron | compound eye photoreceptor development | 46 |
| GO:0042063 | neuron | gliogenesis | 37 |
| GO:0042074 | morphology | cell migration involved in gastrulation | 60 |
| GO:0042127 | morphology | regulation of cell proliferation | 276 |
| GO:0042332 | signalling | gravitaxis | 6 |
| GO:0042673 | neuron | regulation of retinal cone cell fate specification | 82 |
| GO:0042676 | neuron | compound eye cone cell fate commitment | 82 |
| GO:0042745 | morphology | circadian sleep/wake cycle | 6 |
| GO:0042749 | morphology | regulation of circadian sleep/wake cycle | 6 |
| GO:0042803 | signalling | protein homodimerization activity | 155, 32, 37 |
| GO:0042981 | signalling | regulation of apoptotic process | 134 |
| GO:0043005 | neuron | neuron projection | 6 |
| GO:0043053 | signalling | dauer entry | 113 |
| GO:0043066 | signalling | negative regulation of apoptotic process | 32 |
| GO:0043086 | signalling | negative regulation of catalytic activity | 153 |
| GO:0043152 | signalling | induction of bacterial agglutination | 2 |
| GO:0043282 | muscle | pharyngeal muscle development | 71 |
| GO:0043401 | signalling | steroid hormone mediated signaling pathway | 114 |
| GO:0043524 | neuron | negative regulation of neuron apoptotic process | 51 |
| GO:0043620 | regulation | regulation of DNA-dependent transcription in response to stress | 60 |
| GO:0045087 | signalling | innate immune response | 233 |
| GO:0045165 | morphology | cell fate commitment | 37 |
| GO:0045179 | signalling | apical cortex | 82 |
| GO:0045214 | signalling | sarcomere organization | 16, 5 |
| GO:0045468 | neuron | regulation of R8 cell spacing in compound eye | 37 |
| GO:0045475 | neuron | locomotor rhythm | 6 |
| GO:0045500 | neuron | sevenless signaling pathway | 46 |
| GO:0045572 | signalling | positive regulation of imaginal disc growth | 102 |
| GO:0045664 | neuron | regulation of neuron differentiation | 82 |
| GO:0045676 | neuron | regulation of R7 cell differentiation | 82 |
| GO:0045746 | signalling | negative regulation of Notch signaling pathway | 32 |
| GO:0045793 | signalling | positive regulation of cell size | 120 |
| GO:0045892 | regulation | negative regulation of transcription, DNA-dependent | 37 |
| GO:0045893 | regulation | positive regulation of transcription, DNA-dependent | 102 |
| GO:0045900 | regulation | negative regulation of translational elongation | 285 |
| GO:0045944 | signalling | positive regulation of transcription from RNA polymerase II promoter | 101, 114, 155 |
| GO:0045980 | signalling | negative regulation of nucleotide metabolic process | 153 |
| GO:0046008 | signalling | regulation of female receptivity, post-mating | 2 |
| GO:0046331 | signalling | lateral inhibition | 18 |
| GO:0046488 | signalling | phosphatidylinositol metabolic process | 14, 2 |
| GO:0046692 | signalling | sperm competition | 2 |
| GO:0046716 | muscle | muscle cell homeostasis | 16, 5 |
| GO:0046843 | morphology | dorsal appendage formation | 32 |
| GO:0046854 | signalling | phosphatidylinositol phosphorylation | 14, 2 |
| GO:0046928 | neuron | regulation of neurotransmitter secretion | 31 |
| GO:0046982 | signalling | protein heterodimerization activity | 101, 37 |
| GO:0046983 | signalling | protein dimerization activity | 2 |
| GO:0047291 | signalling | lactosylceramide alpha-2, 3-sialyltransferase activity | 3 |
| GO:0048102 | signalling | autophagic cell death | 323, 32 |
| GO:0048190 | morphology | wing disc dorsal/ventral pattern formation | 323, 37 |
| GO:0048311 | regulation | mitochondrion distribution | 54 |
| GO:0048368 | morphology | lateral mesoderm development | 3 |
| GO:0048477 | morphology | oogenesis | 114, 32 |
| GO:0048488 | neuron | synaptic vesicle endocytosis | 134, 46 |
| GO:0048500 | signalling | signal recognition particle | 285 |
| GO:0048512 | morphology | circadian behavior | 6 |
| GO:0048514 | morphology | blood vessel morphogenesis | 63 |
| GO:0048644 | muscle | muscle organ morphogenesis | 5 |

... Table S4 continued
| GO-ID | classification | GO description | internal COP-ID (ake-<br>###) |
| --- | --- | --- | --- |
| GO:0048666 | neuron | neuron development | 37 |
| GO:0048676 | neuron | axon extension involved in development | 203 |
| GO:0048738 | muscle | cardiac muscle tissue development | 16 |
| GO:0048741 | muscle | skeletal muscle fiber development | 71 |
| GO:0048749 | neuron | compound eye development | 32, 37, 51 |
| GO:0048752 | morphology | semicircular canal morphogenesis | 174 |
| GO:0048755 | neuron | branching morphogenesis of a nerve | 3 |
| GO:0048812 | neuron | neuron projection morphogenesis | 3 |
| GO:0048813 | neuron | dendrite morphogenesis | 37, 51, 82 |
| GO:0048844 | morphology | artery morphogenesis | 6 |
| GO:0048846 | neuron | axon extension involved in axon guidance | 14 |
| GO:0048854 | neuron | brain morphogenesis | 37, 55 |
| GO:0048885 | neuron | neuromast deposition | 82 |
| GO:0048886 | neuron | neuromast hair cell differentiation | 82 |
| GO:0048920 | neuron | posterior lateral line neuromast primordium mi-<br>gration | 162 |
| GO:0050768 | neuron | negative regulation of neurogenesis | 37 |
| GO:0050771 | signalling | negative regulation of axonogenesis | 82 |
| GO:0050790 | signalling | regulation of catalytic activity | 3, 63 |
| GO:0050830 | signalling | defense response to Gram-positive bacterium | 153 |
| GO:0050909 | signalling | sensory perception of taste | 82 |
| GO:0050914 | signalling | sensory perception of salty taste | 3 |
| GO:0051124 | neuron | synaptic growth at neuromuscular junction | 31 |
| GO:0051216 | morphology | cartilage development | 174, 3 |
| GO:0051260 | signalling | protein homooligomerization | 3 |
| GO:0051403 | signalling | stress-activated MAPK cascade | 155 |
| GO:0051608 | signalling | histamine transport | 64 |
| GO:0055002 | muscle | striated muscle cell development | 71 |
| GO:0055059 | neuron | asymmetric neuroblast division | 82 |
| GO:0055060 | neuron | asymmetric neuroblast division resulting in gan-<br>glion mother cell formation | 82 |
| GO:0060028 | neuron | convergent extension involved in axis elongation | 400, 60 |
| GO:0060029 | morphology | convergent extension involved in organogenesis | 60 |
| GO:0060037 | morphology | pharyngeal system development | 174 |
| GO:0060041 | neuron | retina development in camera-type eye | 64 |
| GO:0060052 | neuron | neurofilament cytoskeleton organization | 51 |
| GO:0060385 | neuron | axonogenesis involved in innervation | 82 |
| GO:0060438 | morphology | trachea development | 174 |
| GO:0060857 | neuron | establishment of glial blood-brain barrier | 152 |
| GO:0060971 | morphology | embryonic heart tube left/right pattern formation | 3 |
| GO:0061327 | morphology | anterior Malpighian tubule development | 174 |
| GO:0070278 | signalling | extracellular matrix constituent secretion | 174 |
| GO:0070527 | signalling | platelet aggregation | 3 |
| GO:0070829 | signalling | heterochromatin maintenance | 155 |
| GO:0070983 | neuron | dendrite guidance | 82 |
| GO:0071436 | signalling | sodium ion export | 24, 3 |
| GO:0071470 | signalling | cellular response to osmotic stress | 155 |
| GO:0071688 | muscle | striated muscle myosin thick filament assembly | 203, 5 |
| GO:0071805 | signalling | potassium ion transmembrane transport | 144, 26 |
| GO:0072320 | signalling | volume-sensitive chloride channel activity | 19 |
| GO:0072583 | signalling | clathrin-mediated endocytosis | 46 |
| GO:0090305 | signalling | nucleic acid phosphodiester bond hydrolysis | 131 |
| GO:2000223 | signalling | regulation of BMP signaling pathway involved in<br>heart jogging | 3 |

**Table S5.**
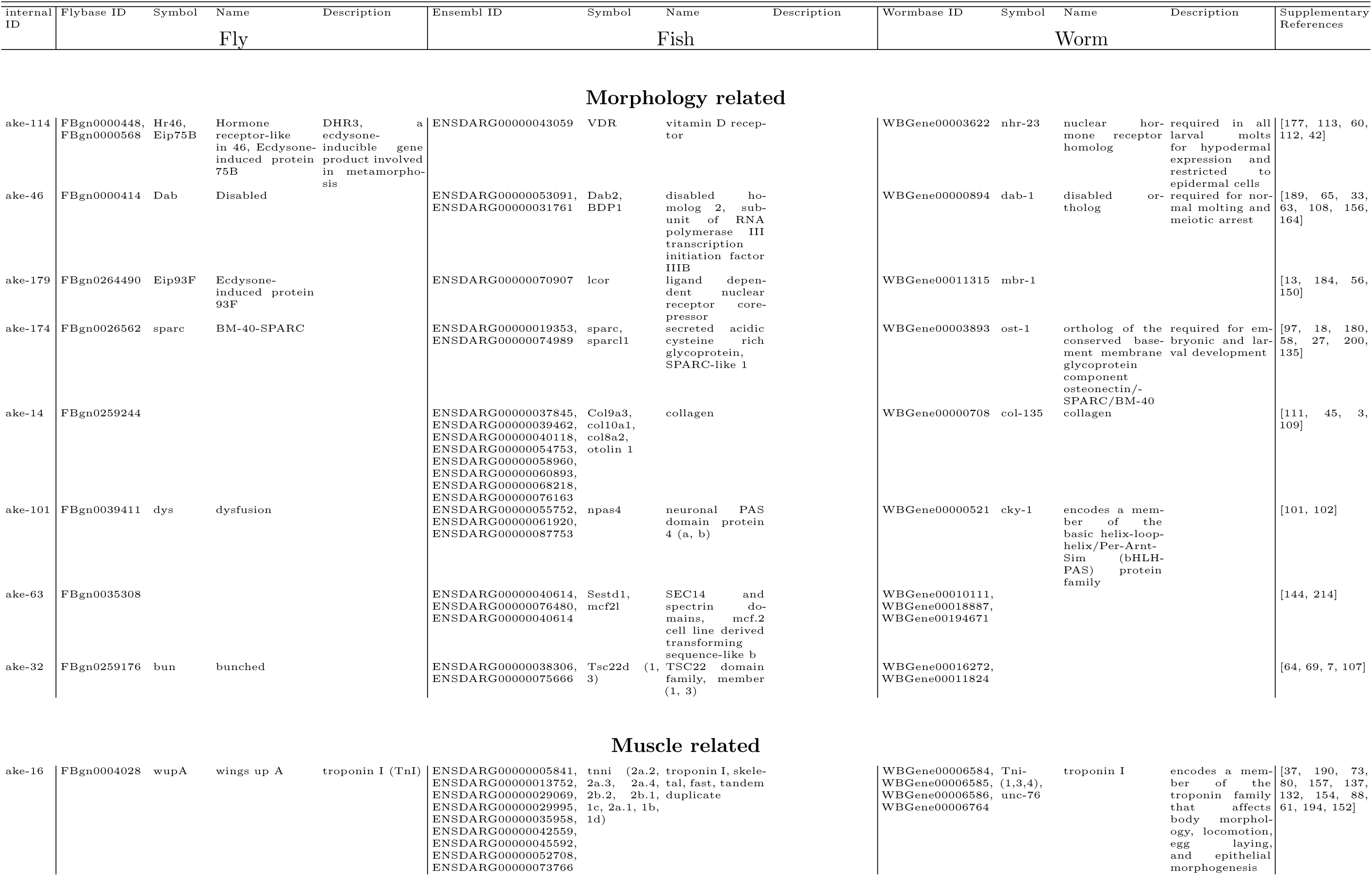
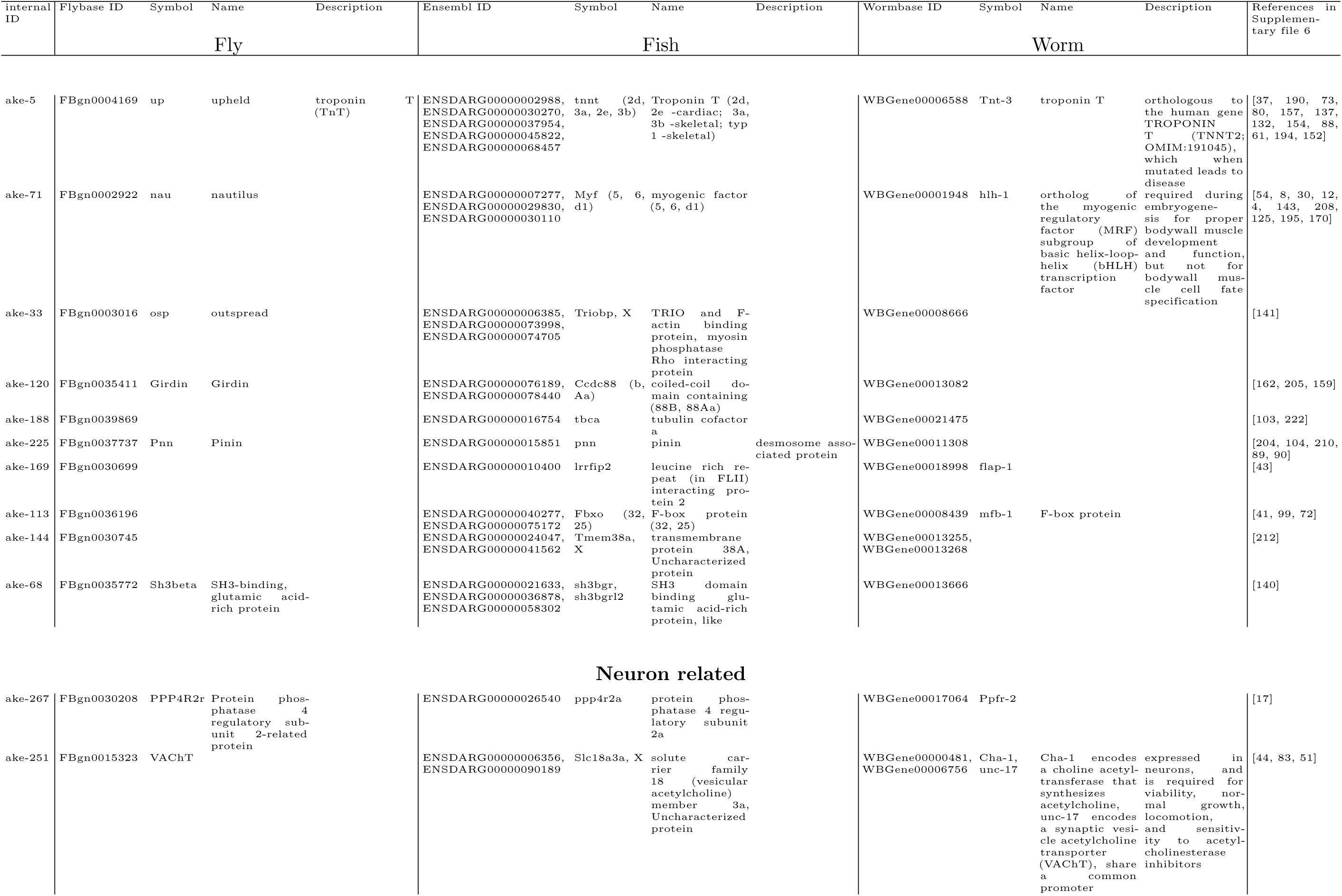
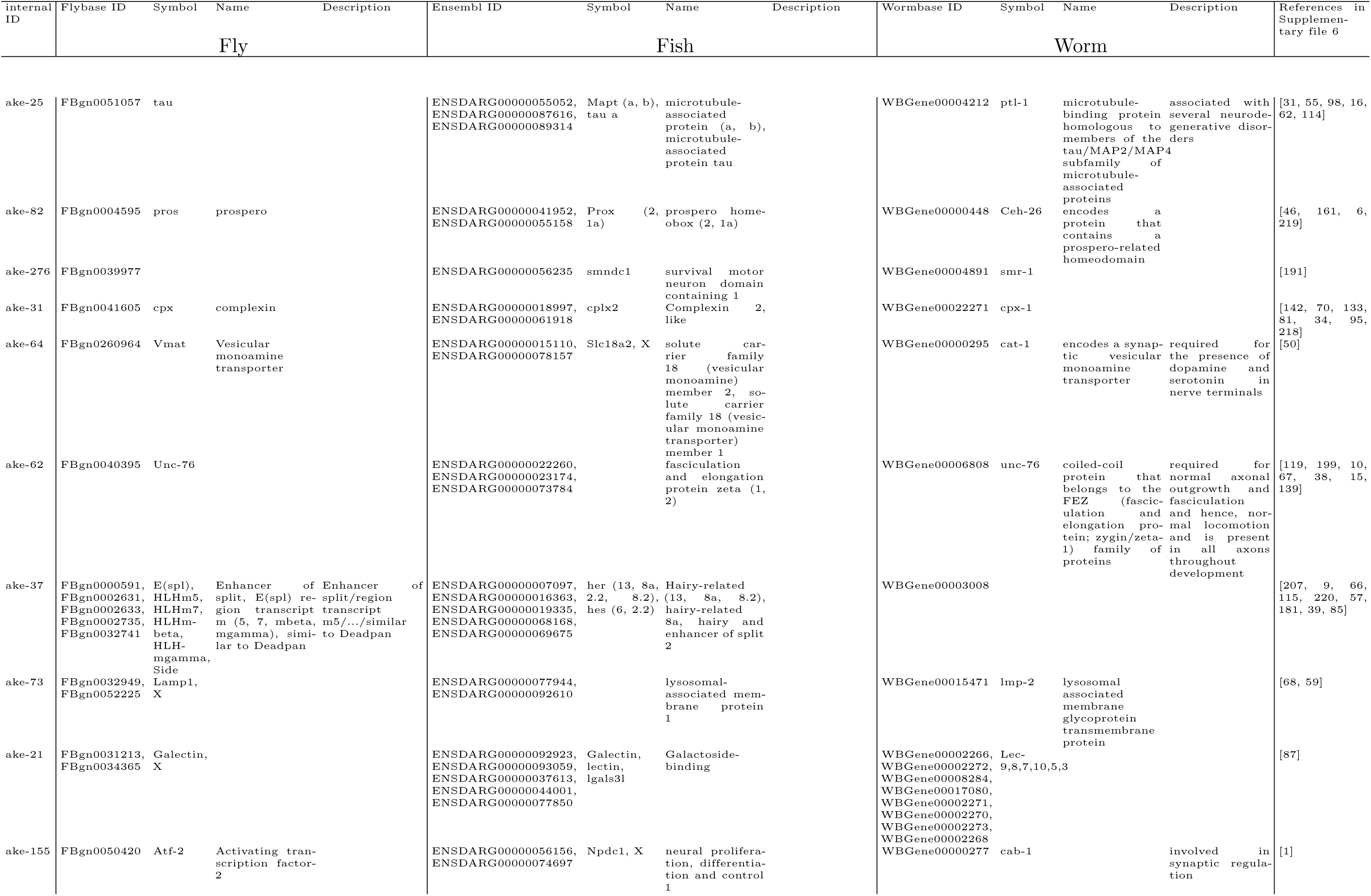
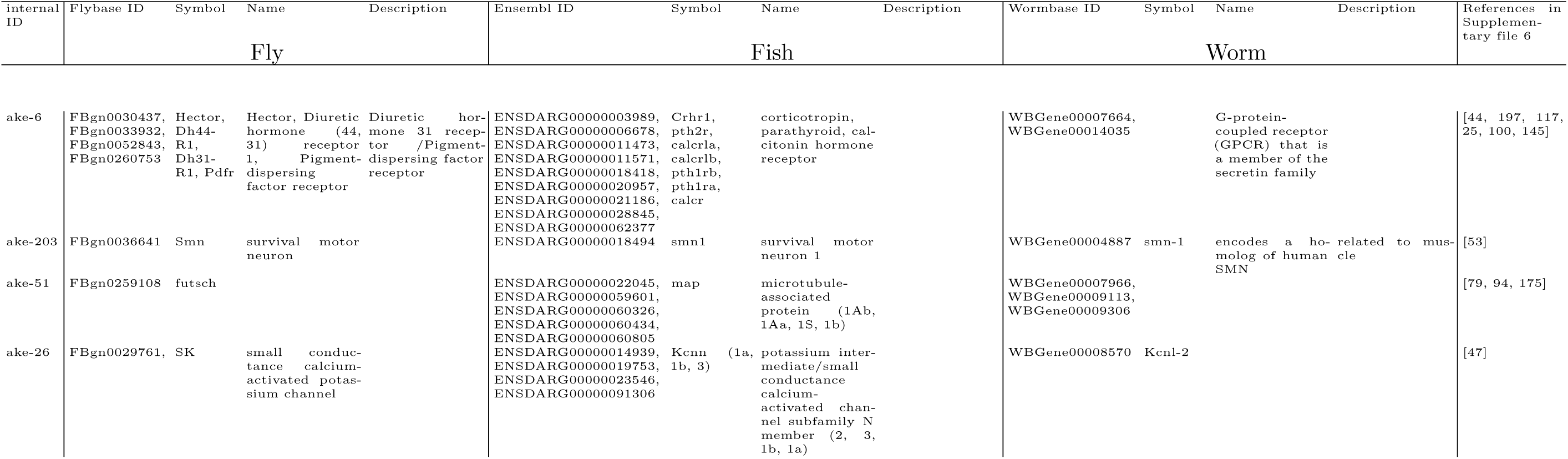
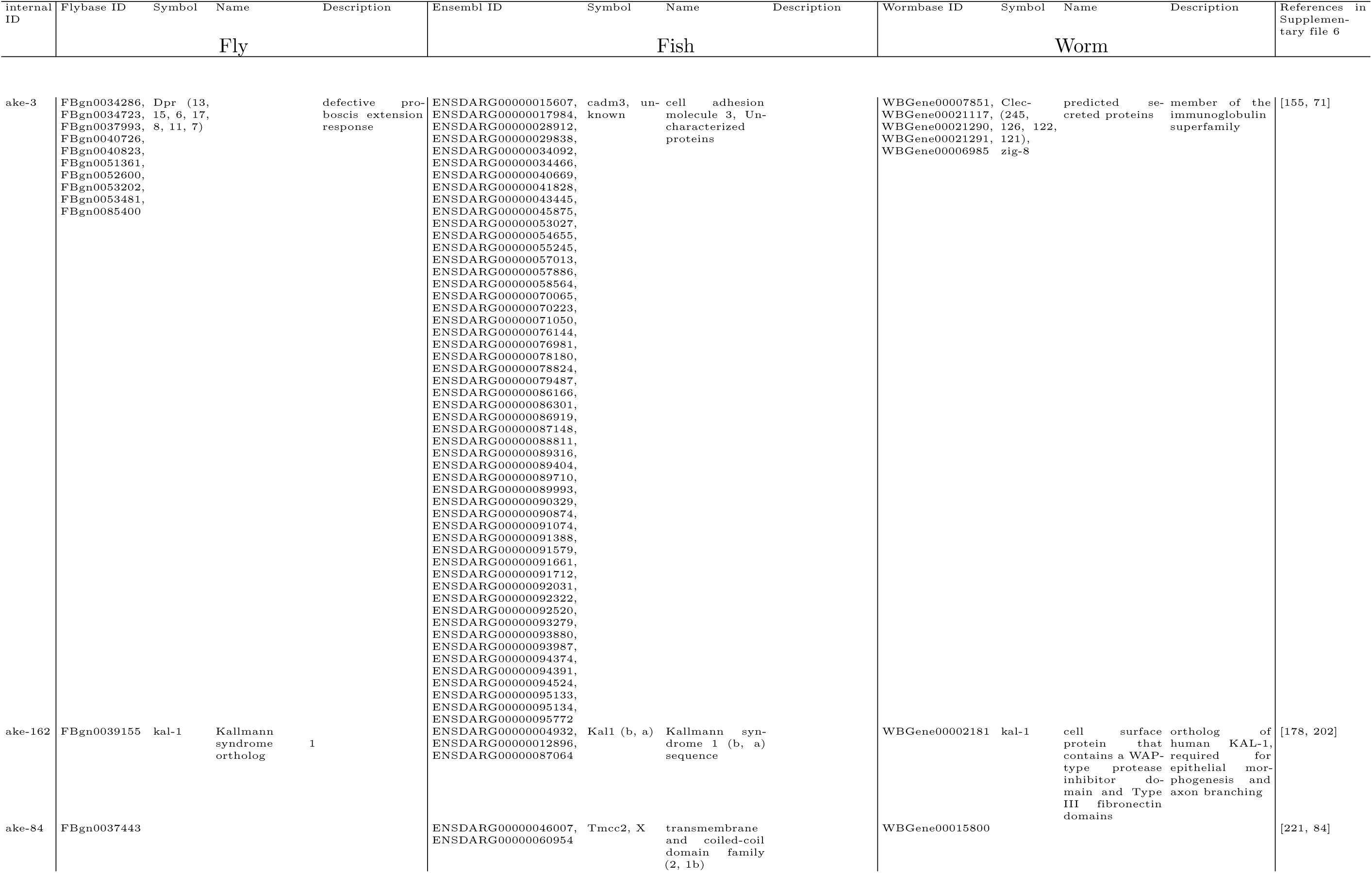
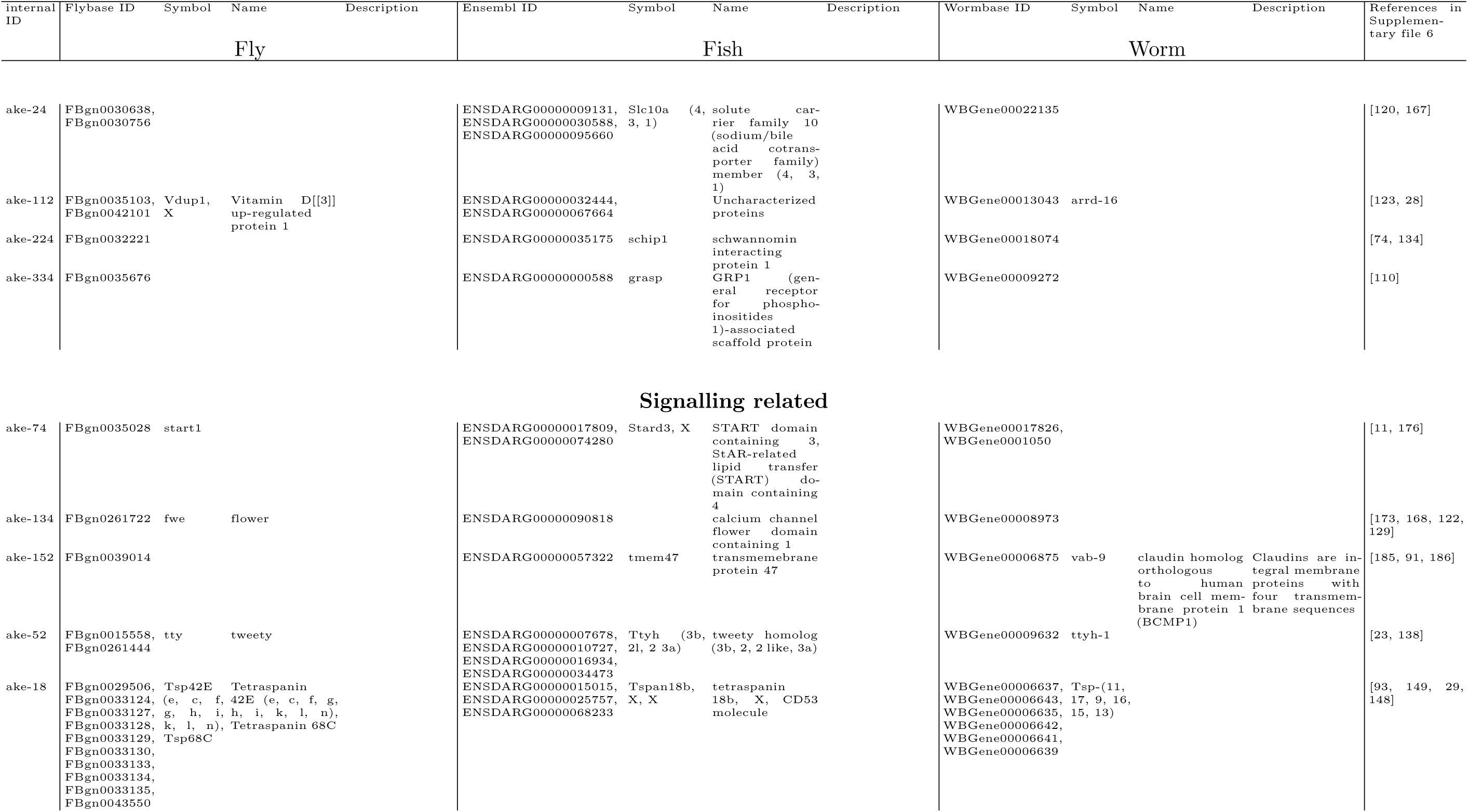
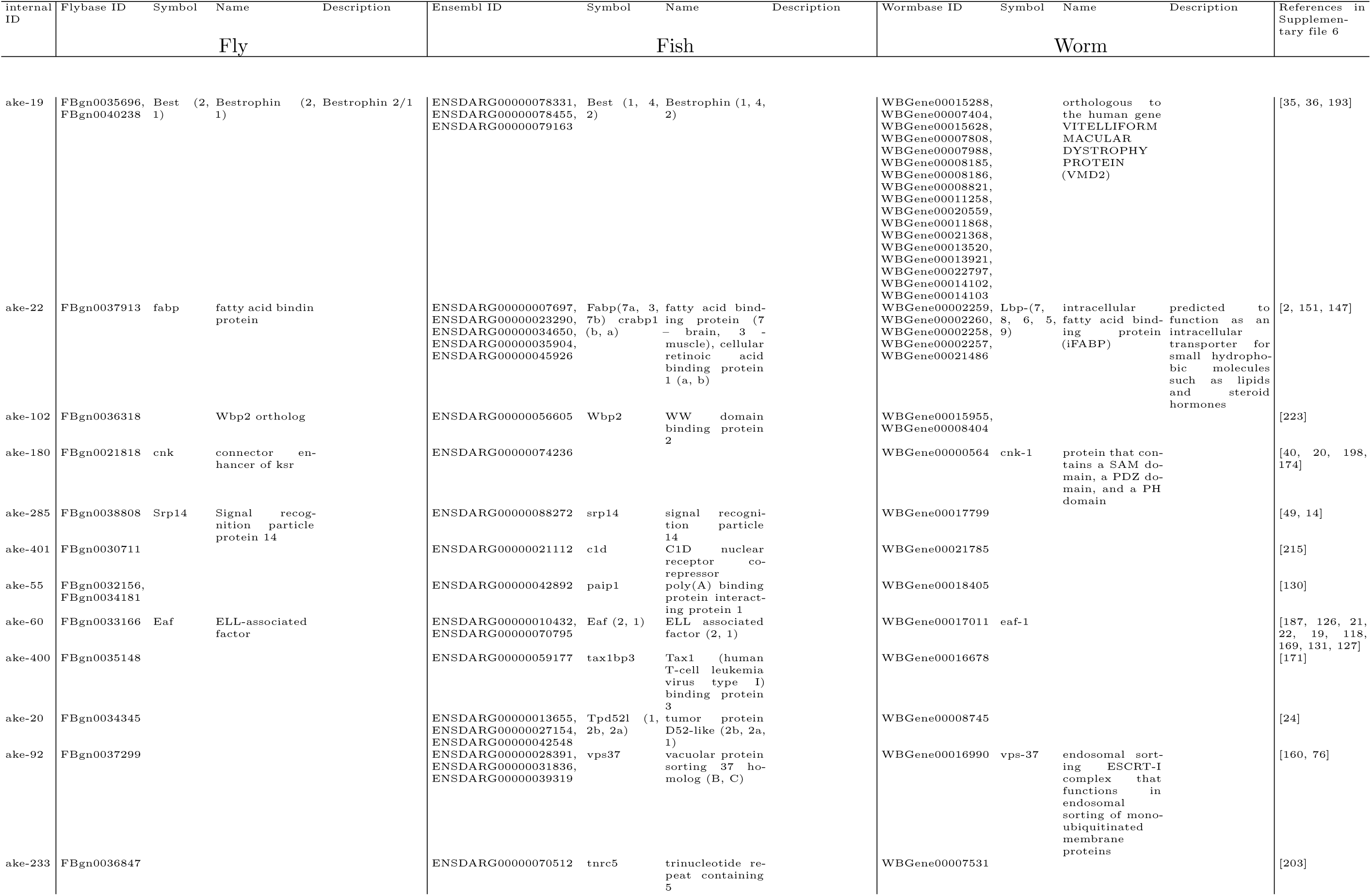
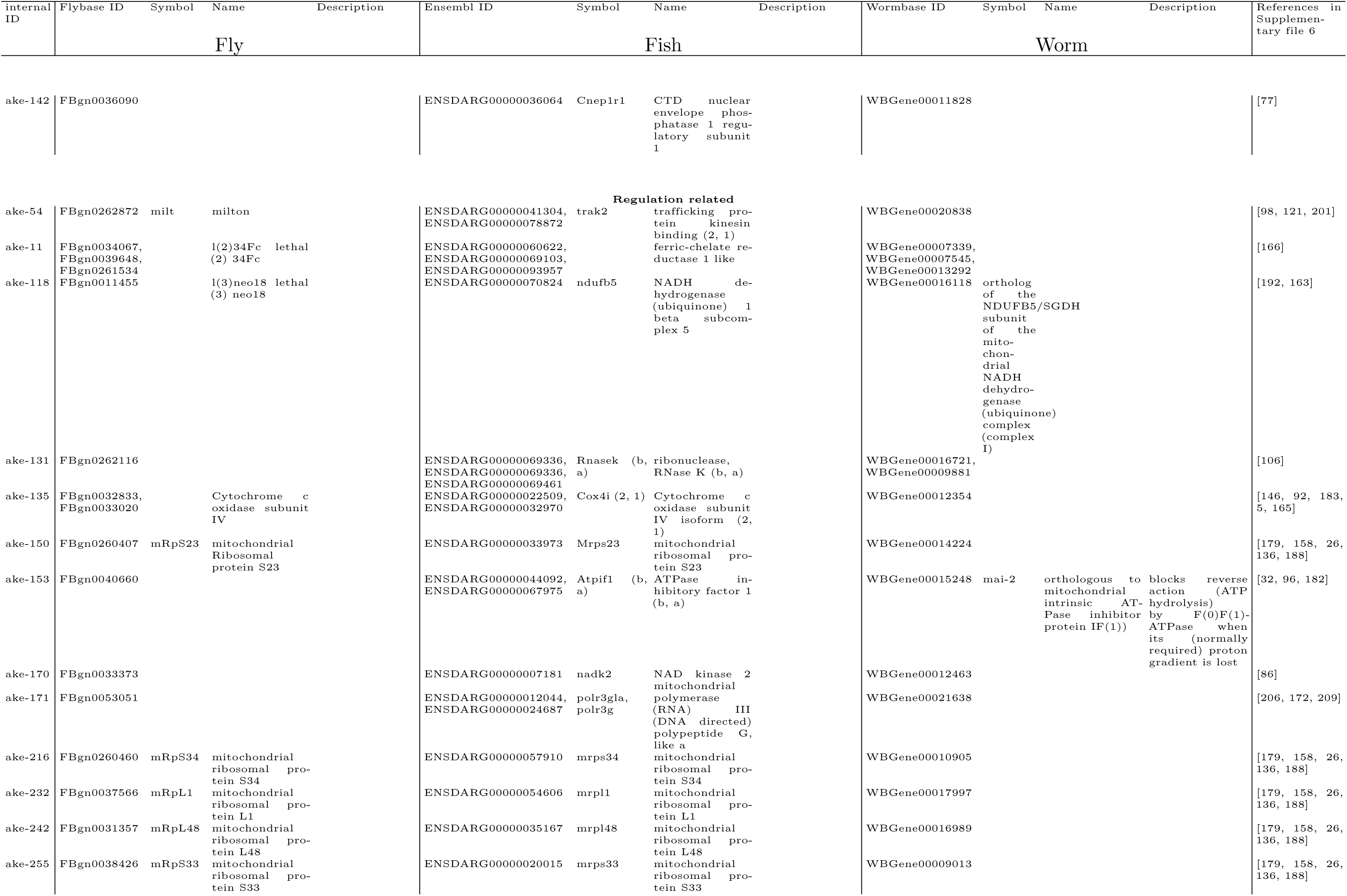
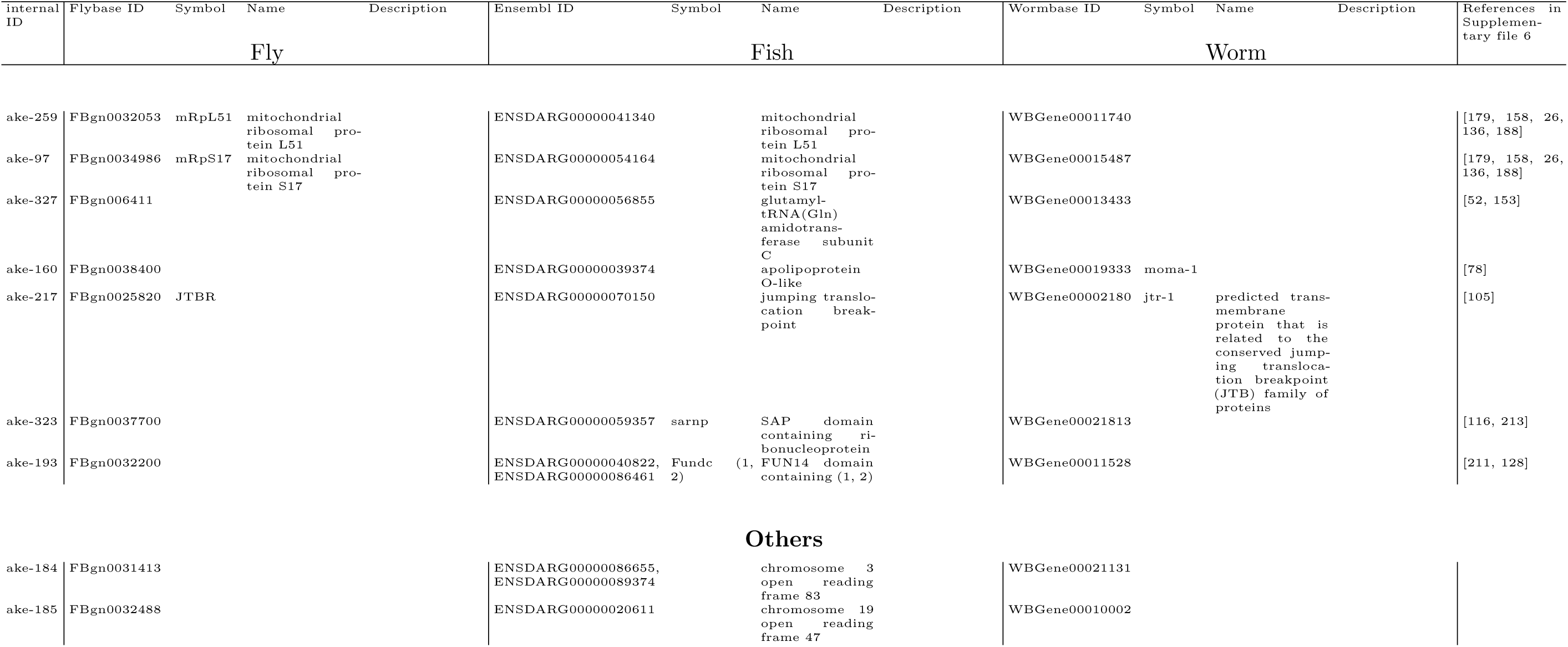
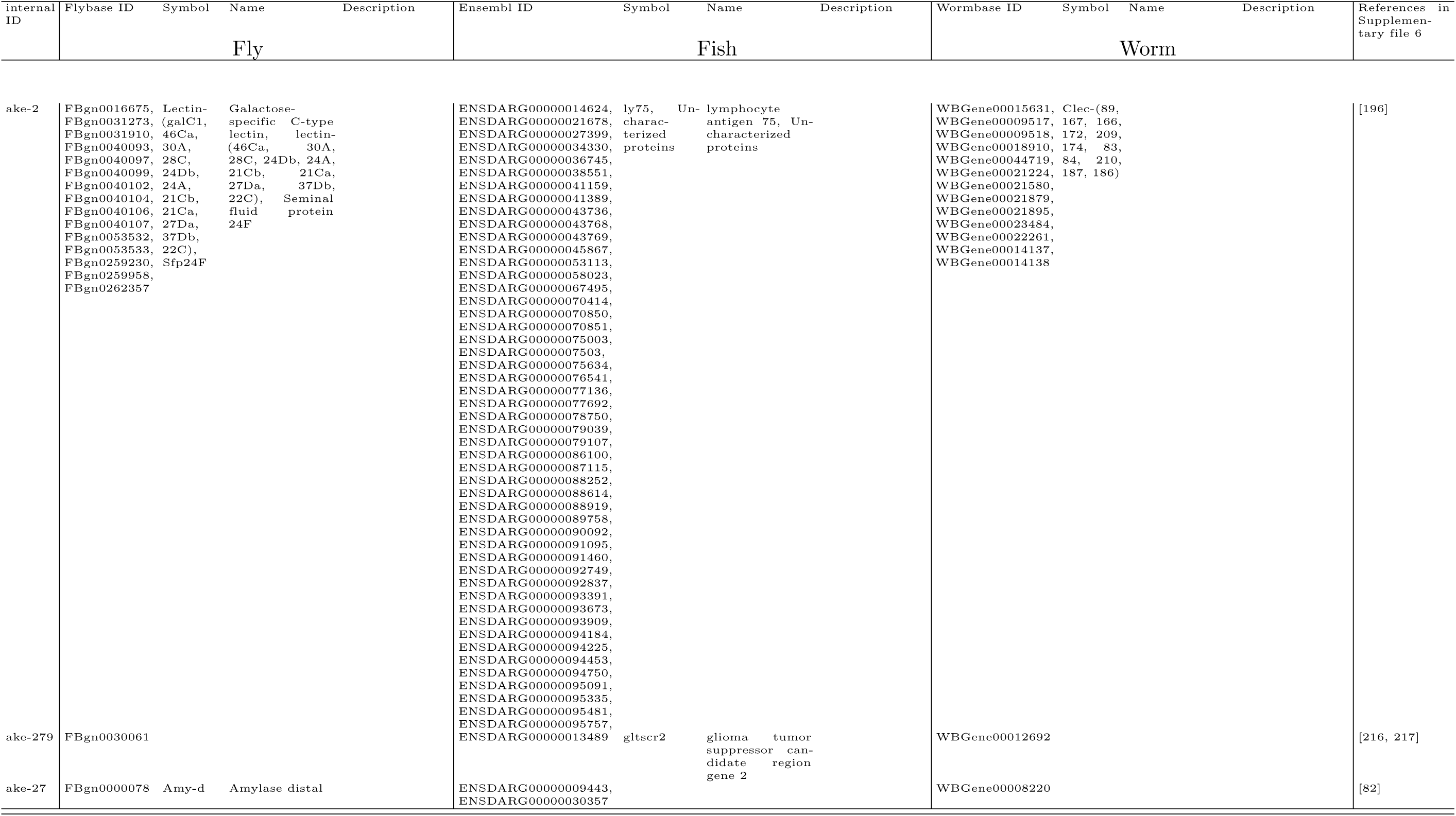
Supplementary Table related to Figures 1 and 4. Description of the 85 clusters in set *L*′.

